# The natural selection of metabolism and mass selects allometric transitions from prokaryotes to mammals

**DOI:** 10.1101/084624

**Authors:** Lars Witting

## Abstract

Inter-specific body mass allometries can evolve from the natural selection of mass, with ±1/4 and ±3/4 exponents following from the geometry of intra-specific interactions when density dependent foraging occurs in two spatial dimensions (2D, Witting, 1995). The corresponding values for three dimensional interactions (3D) are ±1/6 and ±5/6.

But the allometric exponents in mobile organisms are more diverse than the prediction. The exponent for mass specific metabolism tends to cluster around −1/4 and −1/6 in terrestrial and pelagic vertebrates, but it is strongly positive in prokaryotes with an apparent value around 5/6 (DeLong et al., 2010). And a value around zero has been reported in protozoa, and on the macro evolutionary scale from prokaryotes over larger unicells to multicellular vertebrates (Makarieva et al., 2005, 2008).

I show that mass specific metabolism can be selected as the pace of the resource handling that generates net energy for self-replication and the selection of mass, and that this selection of metabolism and mass is sufficient to explain metabolic exponents that decline from 5/6 over zero to −1/6 in 3D, and from 3/4 over zero to −1/4 in 2D. The decline follows from a decline in the importance of mass specific metabolism for the selection of mass, and it suggests *i*) that the body mass variation in prokaryotes is selected from primary variation in mass specific metabolism, *ii*) that the variation in multicellular animals are selected from primary variation in the handling and/or densities of the underlying resources, *iii*) that protozoa are selected as an intermediate lifeform between prokaryotes and multicellular animals, and *iv*) that macro evolution proceeds along an upper bound on mass specific metabolism.

## 1 Introduction

It is difficult to overestimate the importance of body mass allometries in evolutionary biology as they reveal the joint evolution of the life history with mass across the tree of life.

The most well-known allometry is Kleiber (1932) scaling in multicellular animals with a negative 1/4 exponents for the dependence of mass specific metabolism on mass. Yet, the real value, or rather values, of the exponent is still being debated (see e.g., McNab 2008; White et al. 2009; Isaac and Carbone 2010), and it is also questioned whether the functional relationship between the two traits is a straight allometric line or a convexly bend curve (Kolokotrones et al. 2010; Deeds 2011; Ehnes et al. 2011; MacKay 2011).

A value about –1/4 is though in general agreement with the average exponent for the basal (BMR) and field (FMR) metabolic rates across a wide range of vertebrates (e.g., Peters 1983; Savage et al. 2004; Glazier 2005; Duncan et al. 2007; Kabat et al. 2008; Capellini et al. 2010), and the exponents for BMR and FMR are statistically indistinguishable in most lineages of mammals (Capellini et al. 2010). It is therefore reasonable to assume that at a value that is relatively close to −1/4 is acting as a common attractor for the natural selection of allometric relationships, and that this value can vary with variation in the underlying mechanism of natural selection.

It is also essential to recall that the metabolic allometry is only one of several essential allometries, with the empirical exponents in many studies approximating 1/4 for lifespan and reproductive periods, –1/4 for the rate of exponential population growth, –3/4 for the density of populations, and 1 for the area of the home range (Bonner 1965; Schoener 1968; Turner et al. 1969; Fenchel 1974; Damuth 1981, 1987; Peters 1983; Calder 1984).

The correlation between mass and metabolism is not only the most studied allometry empirically; it is also the relationship that has received most theoretical interest (reviewed by e.g. Glazier 2005; White and Kearney 2013). The widespread view has seen the metabolic allometry as a consequence of the physiological resource transportation systems in organisms (e.g. West et al. 1997, 1999a,b; Banavar et al. 1999; Dodds et al. 2001; Dreyer and Puzio 2001; Rau 2002; Santillán 2003; Glazier 2010). Yet, this hypothesis is so far inferior in the sense that it is not integrated with the natural selection of the metabolism and mass that is a pre-condition for the evolution of large organisms with metabolic allometries. The most parsimonious evolutionary explanation is instead the density dependent foraging model of Witting (1995, 1997), where the values of the allometric exponents are selected directly by the natural selection that is necessary for the evolution of large body masses.

The latter model explains all of the above-mentioned exponents from the geometrical constraints of an ecological foraging process that is optimized for a trade-off between the local resource exploitation of the individual and the density dependent interactive competition between the individuals in the population. This interference competition is also the essential factor that selects net assimilated energy into non-negligible body masses in species where the individuals have a sufficiently high net assimilation of resources from the environment (see review by Witting 2008).

With the predicted exponents following from the ecological geometry of foraging, their values are dependent on the spatial dimensionality (*d*) of the foraging process, with 1/4 being the two dimensional case of the more general 1/2*d*. The 1/4 value is therefore predicted to be replaced by 1/6 across species that forage in three spatial dimensions, with a 1/4 → 1/6 like transition being observed quite commonly between terrestrial and pelagic animals (Witting 1995, 1997).

The observed metabolic exponent is though more diverse than the predictions of the ecological model. The empirical exponent varies at least to some degree with mass (Kolokotrones et al. 2010), among major taxa and phylogenetic lineages (Peters 1983; Glazier 2005; Duncan et al. 2007; White et al. 2007, 2009; Sieg et al. 2009; Capellini et al. 2010), and it is also dependent on the activity level of individuals (Darveau et al. 2002; Weibel et al. 2004; Glazier 2005, 2008, 2009; Niven and Scharlemann 2005; White et al. 2007).

More recent studies have also found that the exponent tends to change across the tree of life. Instead of being negative, it is strongly positive in prokaryotes with an apparent value around 5/6 (DeLong et al. 2010), and it has been reported to be zero in protozoa (Makarieva et al. 2008; DeLong et al. 2010) and on the macro evolutionary scale across all non-sessile organisms (Makarieva et al. 2005, 2008; Kiørboe and Hirst 2014).

This variation is not explained by Witting’s (1995, 1997) model, where the selection of mass is independent of the selection on mass specific metabolism. I do therefore in this paper extend the ecological model with primary selection on mass specific metabolism, in order to examine if the joint selection of metabolism and mass will explain the wider set of allometries that is observed across the three of life.

## 2 Basic selection relations

The proposed model is developed to explain average exponents as the evolutionary consequence of a base-case type of interactive ecology between the individuals in a population. Deviations in the ecology from the basecase may result in the evolution of alternative allometries, but this is not studied directly in this paper.

The major difference from my original allometric model (Witting 1995, 1997) is the inclusion of primary selection on metabolism. This implies a mass specific metabolism that is selected as the pace of the resource handling that generates net energy for self-replication, with the selection of mass and allometries following from a predicted increase in net energy. This evolution of metabolism, mass and allometries follows from three basic principles: *i*) the conservation of energy, *ii*) the demography of age-structure, and *iii*) the unfolding of density dependent interactive competition from the population growth of self-replication.

Life history traits within and between natural species tend to correlate in allometric relations, and I use the interactive selection that unfolds from the population growth of self-replication to explain the evolved correlations from uncorrelated traits. This is done by a deterministic model with no contingency (Witting 1997, 2008), which implies that I will explain the evolution of all the life history traits of my model organism. And with a model that is based on the population dynamic feed-back selection of interactive competition, I will also explain ecological traits like home range and abundance.

I will though not attempt to explain absolute trait values, but only the selection response of the life history and the ecology to primary selection on mass specific metabolism and mass. This is done by two processes that I refer to as the metabolic-rescaling and mass-rescaling selection of the life history; with metabolic-rescaling being associated with the pre-mass selection of metabolism that generates net energy for the selection of mass, and mass-rescaling being the selection response of the life history to evolutionary changes in mass. These selection responses are described by the first partial derivatives of the evolving life history with respect to the selected changes in metabolism and mass; with the integrals over mass being the inter-specific body mass allometries.

This level of explanation resembles the Newtonian tradition in physics, where we can explain the acceleration of an object, but not its absolute speed, from the action of a force. In order to use the proposed model to “predict” e.g. an absolute rate of reproduction, I will have to include an observed survival rate as an assumption. The explanation is then no longer deterministic, but contingent upon the observed life history.

To show that the allometric exponents follow from the most basic constraints of the primary selection of metabolism and mass, I will formulate the essential constraints mathematically and solve the resulting selection equations for the unknown exponents. This requires a detailed and consistent description of the essential life history energetics, and of the ecological geometry of density regulation, i.e., of the density dependent foraging and interactive behaviour in one, two and three spatial dimensions.

These descriptions are generally straight forward and they do not include new concepts. But they are nevertheless required to be formulated in a consistent way. For this I will not only refer to my earlier work, but formulate a complete model in the present paper and its appendices, with the essential model parameters being explained in Table 5. I start in Section 2.1 with the new component that is included in the theory, i.e., mass specific metabolism as a primary life history trait. The life history demography and density regulation are described in detail in Appendix A and B, with the relations that are necessary for the allometric deduction being included in Sections 2.2 and 2.3 of the main paper. A short section on the overall relationship between the primary selection of metabolism, net energy and mass is then following, before the deduction of the exponents of body mass allometries in Section 3.

### 2.1 Metabolism, net energy, and time

The metabolic rate is often seen as a proxy for the rate, or pace, at which organisms assimilate, transform and expend energy (reviewed by e.g. Calder 1984; Brown et al. 2004; Humphries and McCann 2014). As such a measure of joint biological activity, metabolism is highly dynamic; being dependent among others on chemistry, temperature, physiology, tissue maintenance and behaviour, with these components being controlled in part by ecological interactions and the age, sexual and informational state of the organism. Yet, for the selection model developed in this paper, all of this variation is integrated into a single measure of the average field metabolic rate per unit body mass (*β*, SI unit J/gs).

Metabolism is transformed into pace by the biochemical, physiological and ecological work that is carried out by mass specific metabolism. Let us therefore define metabolic pace

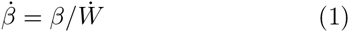

as the frequency (SI unit 1/s) of the mass specific work (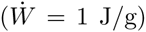) that is carried out by metabolizing one joule per gram of tissue. It is not an absolute given that this work is essential for the organism. If organism metabolism, contrary to our expectations, would evolve by neutral drift instead of by natural selection, the majority of the metabolism would be irrelevant for the ecological and physiological functioning of the organism. But it is fitness costly to burn energy in metabolism. We do therefore expect that the metabolic rate is evolving by a natural selection where metabolic work is used to enhance the fitness of the organism.

To aim for a natural selection explanation, consider the net energy of the organism to be the energy that is available for self-replication. This energy (*∊*, SI unit J/s) is the net energy that is assimilated from a resource per unit time, defined in physical time as a product 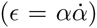 between an ecological/physiological mechanical/biochemical handling of resource assimilation (*α*, resource handling in short, SI unit J), and the pace (α, SI unit 1/s) of this process.

Let us, for a given handling (*α*), assume that the pace of handling 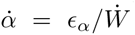 is proportional to the mass specific energy that is used for handling per unit time (*∊_α_*, SI unit J/gs) divided by the mass specific work 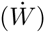 that is carried out by one joule per gram. This energy is provided by metabolism, and we may thus define handling speed by the metabolic fraction 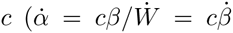, with 0 ≤ *c* ≤ 1) that is used for the handling of net resource assimilation. One, maybe unrealistic, but nevertheless potential example is invariance between handling speed and mass specific metabolism (*c* = *c*_0_/*β*, with 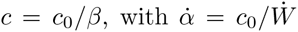 and *c*_0_ being a constant). This can be expected if a fixed amount of energy is used per unit mass on the handling of the resource while the metabolic rate is evolving by other means.

To examine the natural selection of the relationship between metabolism and handling speed, consider net energy as the difference between gross energy (*∊_g_*) and the total energetic cost of the metabolism of the individual (*wβ*):

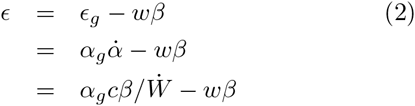

As natural selection favours an increase in *∊* (eqn 20), we find from the positive partial derivative 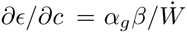 that there is selection for a handling speed that is as large as possible, with the evolutionary equilibrium being *c^**^* = 1 where 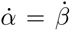. This suggests that mass specific metabolism is selected as a proxy for the pace of resource handling

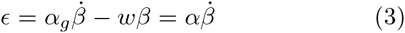

with 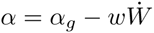 and 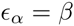.

The energetic difference (*wβ*) between gross and net is then the energy that drives the gross assimilation of energy from the resource. Increasing the speed of handling requires more energy, and this energy is selected to be paid back by a proportional increase in the net energy that is assimilated from the resource. As formulated here, the total metabolic cost (*wβ*) includes not only the energy that is used directly in resource handling, but also the indirect energy of the catabolism and anabolism that goes into the tissue maintenance that is required to keep the organism running.

The net energy

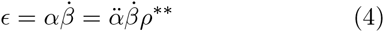

is defined for an implicit resource (*ρ*) at the equilibrium population density (*n^**^*) of the relevant selection attractor on mass, with 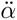 being the intrinsic component of resource handling. Resource handling 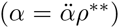 is then specifying the net energy obtained per metabolic pace at the density dependent equilibrium of the selection attractor on mass.

With mass specific metabolism being selected as the pace of the biochemical, physiological and ecological processes of resource assimilation, we can follow Pearl (1928) and others like Brody (1945), Hill (1950), Stahl (1962) and Calder (1984), and consider biological time as inversely related to mass specific metabolism,

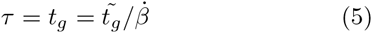

with *τ* being the per generation time-scale of natural selection, *t_g_* the generation time in physical time, and 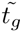 an invariant scaling between generation time and metabolic pace (with SI unit generations/s). The inverse relationship represents an expected invariance when biotic time is defined from metabolic pace; but it is also, as we will see later in Section 3.1, a relationship that is selected directly by the mass-rescaling of the life history.

### 2.2 Life history

The age-structured life history with complete parental investment is described in Appendix A, with the essential energetics that is necessary for the allometric deduction being summarized below.

The first essential component is the total energetic investment in each offspring

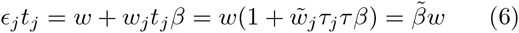

This is the final adult mass *w* of the offspring at the time of independence, plus the energy *w_j_t_j_β* that is metabolized by the offspring during the juvenile period (*t_j_* in physical time; *τ_j_* = *t_j_τ* in biotic time), with *w_j_* being the average juvenile mass during *t_j_* with 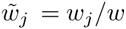, *∊_j_* the energetic investment in the offspring per unit time, and

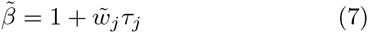

an invariant scaling that accounts for the energy that is metabolized by the offspring. The total investment in each offspring is then reducing to a quantity that is proportional to mass and independent of mass specific metabolism.

This investment implies an energetic quality-quantity trade-off (Smith and Fretwell 1974; Stearns 1992), where the reproductive rate

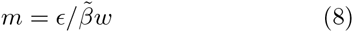

is inversely related to the energetic investment in each offspring. Total lifetime reproduction can then as a consequence (eqn 64) of this be approximated

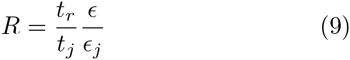

by the product between the ratio of the reproductive period over the juvenile period (*t_r_/t_r_*) and the ratio of net energy over the energy that is allocated to an offspring (*∊/∊_j_*).

The last life history constraint to be considered is the probability that a newborn individual will survive and reproduce

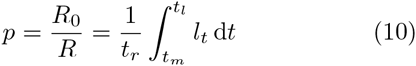

When the *l_t_* function is invariant in biotic time, i.e., when *l_t/t__r_* is invariant, it follows that the 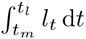 integral reduces to *c_l_t_r_* where *c_l_*, and thus also *p*, are invariant. This is our base-case expectation if survival is determined entirely by intrinsic processes. On top of this there is extrinsic mortality, and if this is about constant in physical time we might expect *p* to be more or less inversely related to *t_r_* with more individuals dying before they reach a given age in biotic time.

### 2.3 Optimal density regulation

The first density regulation constraint that is necessary for the allometric deduction is the adjustment of the life history to an equilibrium (*\**), where the per generation replication (*λ* = *pR*) of the average reproducing unit in the population is one

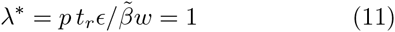

A more essential constraint for the numerical values of the allometric exponents is the ecological geometry of a density regulation that is optimized by natural selection. This was first shown by Witting (1995), and a more general version of his model is described in Appendix B. The essential geometry is that of the density regulated foraging optimum, as it is defined by the spatial dimensions (*d*) of the interactive behaviour and traits like metabolism (*β*), mass (*w*), abundance (*n*), and home range (*h*).

To describe this optimum, I split the overall regulation (*f*) into the three subcomponents

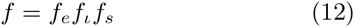

of regulation by exploitative (*f_e_*) and interference (*f_ι_*) competition, and the local exploitation of the resource by foraging self-inhibition (*f_s_*). The latter is not density regulation in itself, but it is needed for the natural selection of a realistic density regulation.

Following the details in Appendix B, in the surroundings of the equilibrium abundance (*n^*^*), the exploitative component can be expressed as

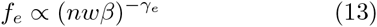

and that of local resource exploitation as

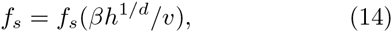

where *f_s_* is a downward bend function that increases monotonically from zero to one as the home range increases from zero to infinity (curve b in Fig. 1), *γ_e_* is the regulation parameter of exploitation, and

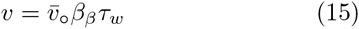

the foraging speed that is defined by the pre-mass component (subscript *β*, see eqn 28) of metabolic pace (given here by its proportional relation with metabolism, *β_β_*), and the mass-rescaling component (subscript *w*) of biotic time (*τ_w_*).

In relation to interference competition, regulation *f_l_* = *e^−lμ^* is a declining function of the level of interference competition (*ι*) and the average cost of interference (*μ*), with the level of interference

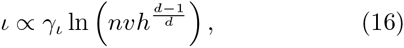

being dependent on the abundance, home range, and foraging speed. Because of the quality-quantity trade-off (eqn 8), where the reproductive rate is increasing with a decline in mass, we can follow eqn 80 and express the level of interference at the population dynamic equilibrium as a declining function of the average mass

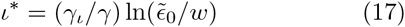

The cost of interference is increasing with an increase in the average home range (Fig. 1, declining curve a), but the cost of foraging self-inhibition is declining (Fig. 1, increasing curve b). Hence, there is an optimal home range, where the joint regulation by interference and self-inhibition is minimal (Fig. 1, c). This optimum

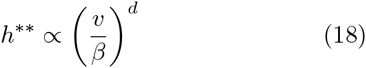

implies that density regulation is body mass invariant as a whole [*f* = *f_e_f_l_f_s_ ∝ w*^0^], and this generates natural selection for a covariance

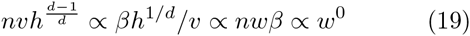

that will leave the regulation optimum, and the level of interference competition in the population (eqn 16), invariant of the life history (see Appendix B.5).

### 2.4 Metabolism, net energy and mass

The next selection that we need to consider for the allometric deduction is primary selection on metabolism, net energy and mass. For this we may combine eqns 4 and 11 and find that the selection gradients

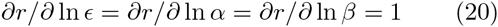

on the log of net energy (*∊*), and its subcomponents *α* and *β*, are unity. The secondary theorem of natural selection (Robertson 1968; Taylor 1996) is therefore predicting an exponential increase

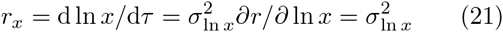

in net energy, resource handling, and mass specific metabolism (*x* ∊ {*∊, α, β_β_*}) on the per generation time-scale (*τ*) of natural selection, when the additive heritable variance 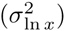 is stable.

**Figure 1:**
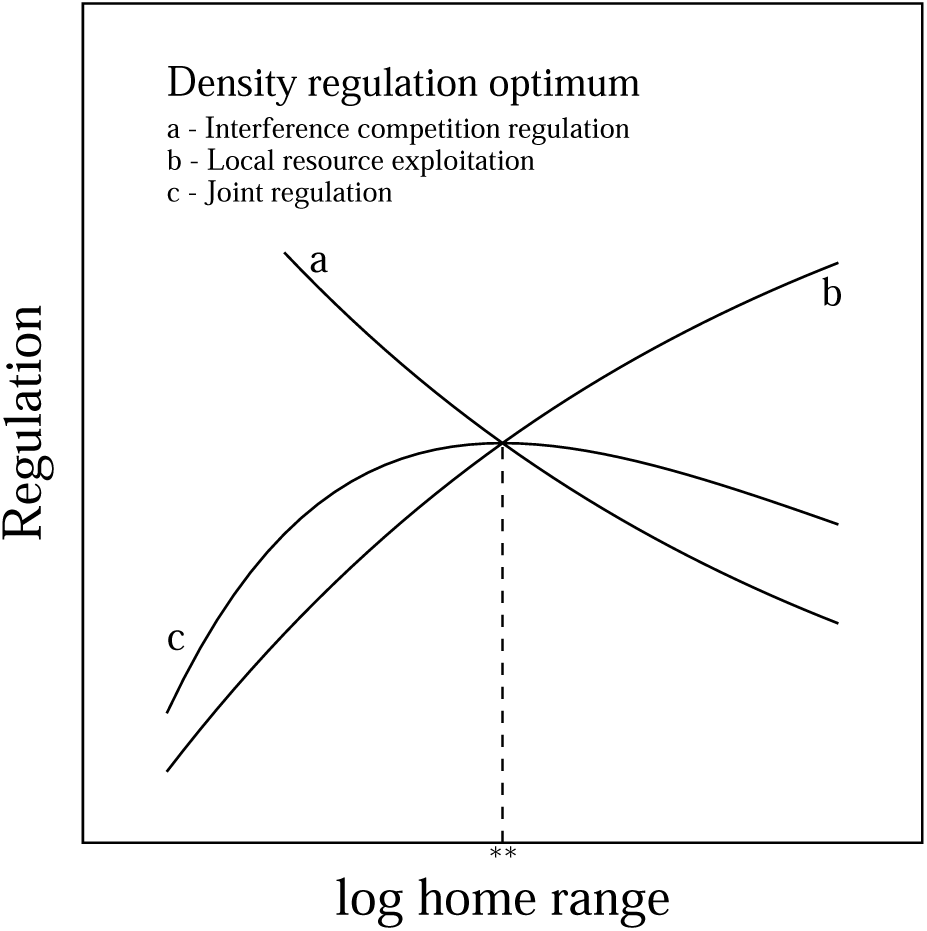
**The optimal home range** (**) as defined by the density regulation optimum for interference competition and local resource exploitation.

While this implies sustained selection for an increase in *∊*, the average net energy may decline due to environmental variation, or inter-specific interactions when a smaller species is excluded from essential resources by larger species. And owing to the competitive exclusion of species by inter-specific competition, we may expect a distribution of species with net energetic states that range from a possible minimum to a maximum, with the maximum increasing over evolutionary time due to eqn 21. I will not directly consider the evolution of the exponential increase in this paper [see Witting (1997, 2003, 2016b,c) instead], but only note that the increase allows me to consider a distribution of species that differ in *α*, *β_β_* and *∊*.

Independently of the selection cause for the evolution of mass, the individuals of an evolutionary lineage cannot be large unless they have evolved the ability to consume plenty of resources. This implies that an evolutionary change in mass is selected, in one way or the other, as a consequence of the evolutionary change in energy

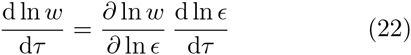

with

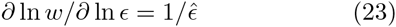

defining the selection dependence of mass on energy given a log-linear relation. The specific mechanisms of this selection are not considered here; they are instead described by Witting (1997, 2008, 2016a,). From eqn 23 we may conclude that natural selection creates an evolutionary function

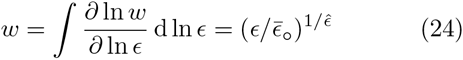

that defines mass as an evolutionary consequence of a selection that is imposed by the net energy of the organism, where 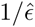 is the allometric exponent that is given by the selection relation of eqn 23, and the intercept 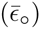 is an initial condition.

## 3 Body mass allometries

The allometries that can be observed across natural species are post-mass allometries in the sense that they are the allometric correlations that have evolved by the complete selection process from the pre-mass selection of net energy, over metabolic-rescaling and body mass selection, to rescaling selection from the evolutionary changes in mass.

The post-mass allometry 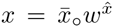 for the dependence of a trait *x* on mass is thus a product

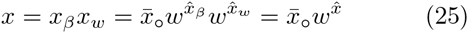

between the post-mass allometry of mass-rescaling

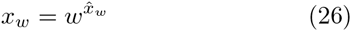

that describes the complete allometric scaling with mass when there is no pre-mass selection on mass specific metabolism, and the post-mass allometry of metabolic-rescaling

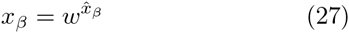

that is the additional allometric scaling that evolves from metabolic-rescaling and the dependence of body mass selection on the net energy that is generated by pre-mass selection on mass specific metabolism (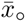 is an unexplained initial condition, and 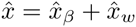).

Many of the allometries that have been established empirically across natural species have Kleiber scaling with exponents around ±1/4 or ±3/4. The life history covariance that evolves from my selection model identifies these exponents as the mass-rescaling of eqn 26. Hence, in some instances we may regard the metabolic-rescaling allometries of eqn 27 as the evolution of the intercepts of the more traditional mass-rescaling allometries; an evolution that is caused by pre-mass selection on metabolic pace.

### 3.1 Mass-rescaling

The inverse of eqn 24 is the traditional allometry 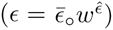 between mass and energy. And with net energy being a function of handling and pace (eqn 4), this implies that the allometric exponents of the three traits *∊*, *α*, and *β* are interrelated. To describe this, let the *∊* ∝ *αβ* relation be rewritten as a product of sub-components of resource handling and mass specific metabolism

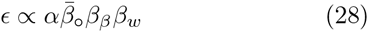

with 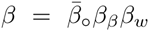 being split into a mass-rescaling 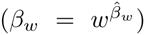 and a pre-mass (*β_β_*) selection component, with 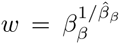 being the selected dependence of mass on the pre-mass component of mass specific metabolism.

Of the three parameters *∊*, *α* and *β* it is only metabolism that is split into a pre-mass and a mass-rescaling component. Resource handling is defined as a pure pre-mass parameter that is generating net energy for the selection of mass independently of the changes in the pace of resource handling. And net energy is found to be invariant of mass-rescaling on the per generation time-scale (see Section 3.1.1) that is relevant for the natural selection of mass.

Relating to the relationship between the allometric exponents we have 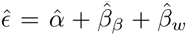 from eqn 28. To identify the essential dependence in this expression, recall that the selection pressure on mass is reflecting the exponent of net energy 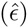, as it is defined by the partial derivative 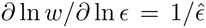 of eqn 24. This implies that we define the selection of mass from the changes in the average net energy invariantly of the underlying causes (*α, β_β_ & ρ*) for the change in energy. We may thus expect that 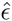 is invariant of the other exponents 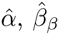 and 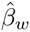, while the latter three are evolutionarily interrelated by the following trade-off

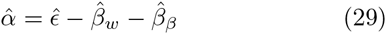

This invariance of the 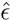 exponent relative to the other three exponents in eqn 29 is confirmed later in this section, together with a similar invariance for the mass-rescaling exponent of metabolism 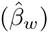. Hence, in the end we will find a direct allometric trade-off between the exponent for resource handling 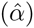 and the pre-mass exponent for mass specific metabolism 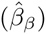. Please note that this trade-off between the 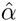 and 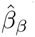 exponent is not reflecting a metabolic-rescaling of resource handling. It is only reflecting the relative importance of resource handling and the pre-mass component of mass specific metabolism for the generation of the variation in the net energy that is responsible for the natural selection of the variation in the body masses of the species that are being compared in an allometric study.

Now let us ignore variation in the pre-mass component of mass specific metabolism, i.e., let

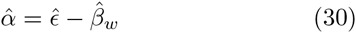

from eqn 29 with 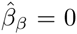 and *x_β_* = *w*^0^ for all traits *x*. This corresponds with the allometric model of Witting (1995), that determines the rescaling of the life history in response to the evolving mass. It will give us allometries for the limit case where all the evolutionary variation in mass is induced by variation in resource handling and/or resource availability, with metabolism evolving exclusively by the allometric rescaling with mass.

In this case, for the other life history and ecological traits (*x*) we can expect an allometric mass-rescaling

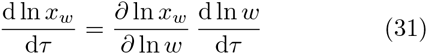

where traits are selected in response to a selection change in mass (eqn 22). Given log-linear selection relations

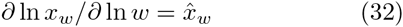

this implies the evolution of mass-rescaling allometries

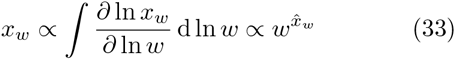

where 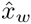 is the mass-rescaling exponent.

#### 3.1.1 Metabolic trade-off selection

Let this mass-rescaling selection be induced by a metabolism that trade-offs against the time that is needed for reproduction, when the parental energy that is allocated to the offspring has to be used either on the growing mass or on the metabolism of the offspring. Less energy will be available per unit time for the mass of the growing offspring when more energy is metabolized, and this will cause the juvenile period to increase, and the reproductive rate to decline, with an increase in the metabolic rate.

The evolutionary linking of the different life history traits by this metabolic trade-off selection is illustrated by the following causal relationships

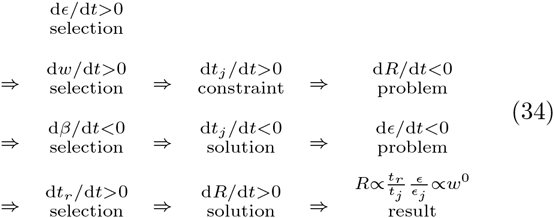

Initially we have the selection induced increase in net energy (d*∊/*d*t >* 0, eqn 21) that allows for a selection increase in mass (d*w/*d*t >* 0, eqn 22). Among the variants with a similar larger than average body mass that is favoured by this selection, it is the variant that has life history traits that are correlated in such a way that the physiological (frequency independent) replication rate is invariant of mass, that is selected over variants where the replication rate is declining with mass. Mass-rescaling by metabolic trade-off selection is the selection of these frequency independent trait correlations that are induced by the more direct selection pressure on mass.

From the physiological constraint of eqn 6 on net energy, metabolism and mass we find that the juvenile period, that defines the time that is needed to grown an offspring 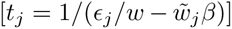, is increasing (d*t_j_*/d*t >* 0) with mass as 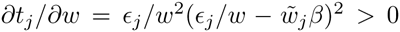. Then, from the reproductive constraint 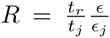 of eqn 9 we have that the increased juvenile period implies a lifetime reproduction and fitness decline (d*R/*d*t <* 0) that is avoided by the selection for increased mass if possible.

This selection conflict is solved in the third line of eqn 34, by trait correlations for an allometric rescaling where mass specific metabolism is declining with mass (dβ/d*t <* 0). This will shorten (d*t_j_*/d*t <* 0) the juvenile period 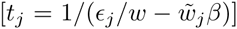 because a larger fraction of the parental energy is then allocated to the growth of the offspring at the cost of the energy that is burned by the metabolism of the offspring. From the total energy invested in an offspring (eqn 6), the expression for lifetime reproduction is 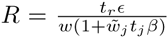. Hence, provided that *t_r_∊* is constant, the selection conflict on mass is cancelled when 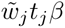 is invariant of mass; implying selection for a mass specific metabolism that is inversely proportional to the juvenile period *β_w_* ∝ 1*/t_j,w_*.

But as 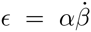, we find that the net energy will decline (d*∊/*d*t <* 0) with the decline in mass specific metabolism, and given 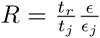 this implies a decline in lifetime reproduction and fitness because the parent will no longer have the required energy available for reproduction. This selection conflict is solved on the last line of eqn 34, by trait correlations that extend the reproductive period (d*t_r_*/d*t >* 0) until it is proportional with the juvenile period and inversely proportional with mass specific metabolism. This leads to an increase in lifetime reproduction (d*R/*d*t >* 0) that results in physiologically invariant fitness

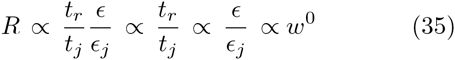

that is selected by the selection for increased mass.

The evolutionary consequence of eqn 35 is a selection that trade-offs (dilates) the per generation time-scale of natural selection in order to maintain net energy and fitness invariant of a selection increase in mass. The result is an energetic state that is maintained constant in biotic time (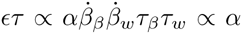, with 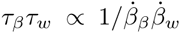) while it is declining in physical time 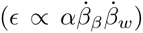 with a decline in mass specific metabolism with mass 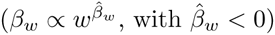expected for time scaling of intrinsic survival.

When the *R* ∝ *w*^0^ invariance is combined with the population dynamic constraint *pR* = 1, we expect a mass invariant survival

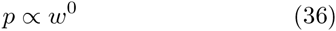

from a survival curve with age that is invariant in biotic time, as expected for time scaling of intrinsic survival (eqn 10). And when the *∊/∊_j_* ∝ *w*^0^ invariance of eqn 35 is combined with the equation for the total energetic investment in each offspring (eqn 6), we obtain the following energetic constraint

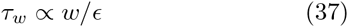

that defines the mass-rescaling component of biotic time from the ratio of the selected mass over the selected net energy.

This mass-rescaling is also including the optimal density regulation of Section 2.3, because the three density regulation components *f_e_*(*nwβ*), 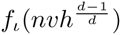 and *f_s_*( *βh*^1*/d*^*/v*) are dependent on the traits that are rescaled by the selection increase in mass. The result is a trait covariance that is selected not only by the constraints of the metabolic trade-off (eqns 35 and 37), but also by the constraints of optimal density regulation (eqn 19). And where it is the metabolic trade-off selection that initiates the selection response to the evolutionary changes in mass it is, as we will see in the next sub-section, primarily the ecological geometry of optimal density regulation that explains the actual values of the allometric exponents.

#### 3.1.2 Allometric deduction

In order to deduce the allometric exponents from the selection conditions that we have already described, let us use *τ* as the scaling parameter for all biotic periods *t_x_* = *τ_x_τ*, and exchange *v* with *τ_w_* from eqn 15. Then, insert power relations 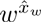 for the different traits *x* ∊ {*p, τ, ∊, β, n, h*} into eqns 5, 37, and 19 to obtain

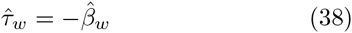

**Table 1:** Allometric rescaling. a) The predicted mass-rescaling exponents 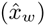 of body mass (*w*) allometries 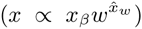 as functions of the spatial dimensions (*d*) of the interactive behaviour. b) The predicted metabolic-rescaling exponents 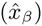 of the mass-rescaling intercepts 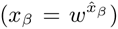 as a function of the metabolic-rescaling exponent for mass specific metabolism 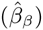. **Symbols:** *∊*:net energy; *α*:resource handling; *β*:mass specific metabolism; *τ*:biotic periods/time; *p*:survival; *R*:lifetime reproduction; *r*:population growth rate; *h*:home range; *n*:population density.

| a) Mass-rescaling as a function of $d$ | | | | | | | | | |
| --- | --- | --- | --- | --- | --- | --- | --- | --- | --- |
| $\hat{\epsilon}$ | $\hat{\alpha}$ | $\hat{\beta}_w$ | $\hat{\tau}_w$ | $\hat{p}_w$ | $\hat{R}_w$ | $\hat{r}_w$ | $\hat{h}_w$ | $\hat{n}_w$ | |
| $\frac{2d-1}{2d}$ | 1 | $-\frac{1}{2d}$ | $\frac{1}{2d}$ | 0 | 0 | $-\frac{1}{2d}$ | 1 | $\frac{1-2d}{2d}$ | |

Table 1: **Allometric rescaling.**
| b) Metabolic-rescaling as a function of $\hat{\beta}_\beta$ | | | | | | | |
| --- | --- | --- | --- | --- | --- | --- | --- |
| $\hat{\alpha}$ | $\hat{\tau}_\beta$ | $\hat{p}_\beta$ | $\hat{R}_\beta$ | $\hat{r}_\beta$ | $\hat{h}_\beta$ | $\hat{n}_\beta$ | |
| $1 - \hat{\beta}_\beta$ | $-\hat{\beta}_\beta$ | $\hat{\beta}_\beta$ | $-\hat{\beta}_\beta$ | $\hat{\beta}_\beta$ | 0 | $-\hat{\beta}_\beta$ | |

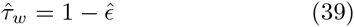

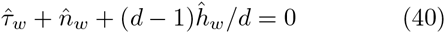

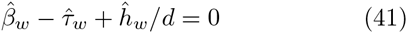

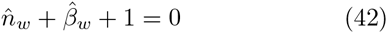

and the following invariance

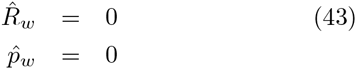

from eqns 35 and 36.

Now, from eqns 42 and 38 we have 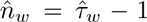. Insert this expression into eqn 40 and obtain 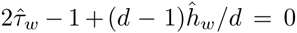, and exchange 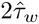 with 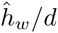 from eqns 41 and 38 to obtain 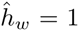. Then, from eqns 41 and 38 we have 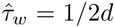 and 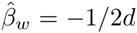; and from eqns 30, 39, and 42 that 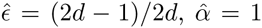 and 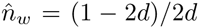. And with the population dynamic growth rate in physical time being *r* = ln(*pR*)*/t_g_* we have 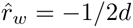 given *pR* ∝ *w*^0^.

These deduced mass-rescaling exponents are listed in Table 1a. They are the same as those of the original deduction in Witting (1995), except that the new deduction is more general as it explains also the exponent for survival (*p*) and the inverse link between the reproductive period and metabolism (*τ* ∝ 1/β).

One way to illustrate the evolutionary significance of the mass-rescaling allometries is to plot the average replication at the population dynamic equilibrium (*pR*) as a function the potentially possible rescaling exponents. This is done in Fig. 2, and it illustrates that it is only for the actual mass-rescaling of the allometric solution that natural selection can maintain the life history in the required balance where the average per generation replication rate at the evolutionarily determined population dynamic equilibrium is one invariantly of mass.

**Figure 2:**
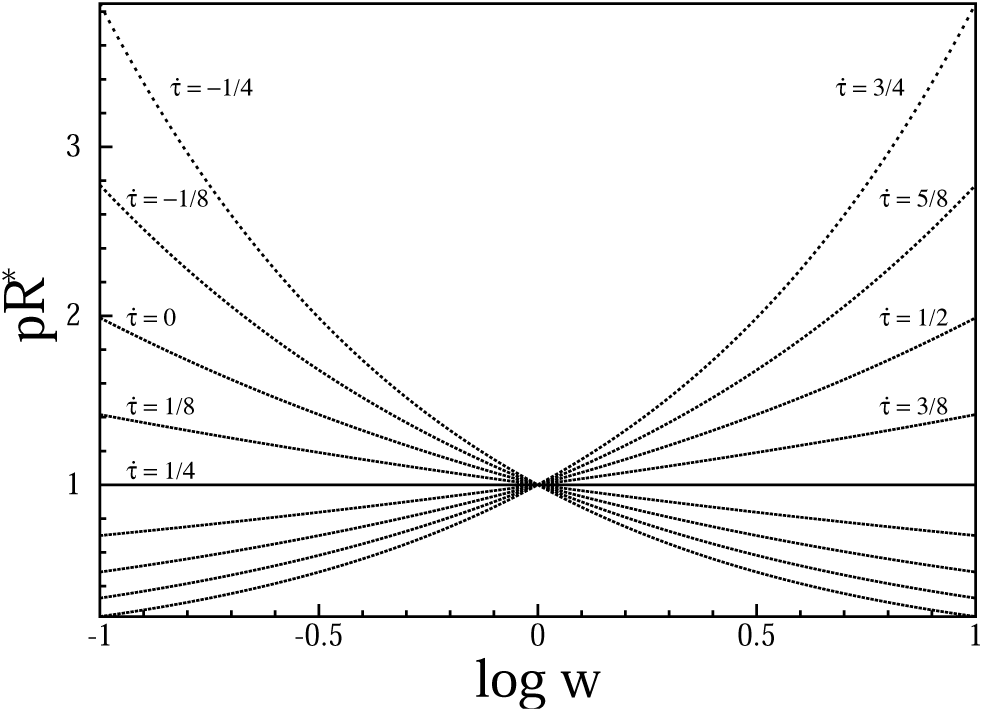
**The replication invariance.** A theoretical span (eqn 96) of the average per generation replication rate (*pR\**) in the population as a function of the selected mass (*w*) for a range of potentially possible mass-rescaling exponents 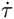. It is only the exponent of the allometric solution 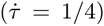 that maintains the life history in the required balance where the average per generation replication rate at the evolutionary optimum is one invariantly of mass. The model behind the figure is given in Appendix C, and the plot is for *z* = 0.1 for eqn 96, given 2D interactions where 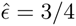.

### 3.2 Metabolic-rescaling

Let us now go beyond the mass-rescaling allometries and consider the influence of variation in the mass-rescaling intercepts as it evolves from pre-mass selection on metabolic pace. With mass specific metabolism (*β*) being selected as the pace 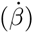 of resource handling (*α*), it is providing part of, or the complete, net energy 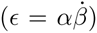 that is driving the evolution of mass, and it is thus influencing the allometric scaling independently of the mass-rescaling of the previous section.

The evolutionary increase in metabolism is causing a metabolic-rescaling

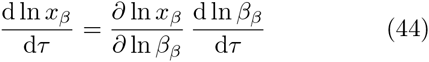

that exists independently of the evolutionary changes in mass. This rescaling affects rate related life history traits that are scaled by the increase in metabolic pace, with subscript *β* denoting components that relate to this pre-mass selection on metabolic pace. This rescaling will transform log-linear selection relations

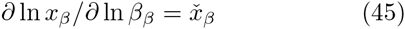

into metabolic allometries

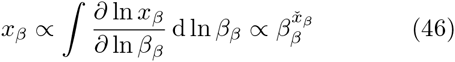

where, among thers, increased mass specific metabolism is shortening biotic time periods like generation time and increasing the amount of resources that the individual assimilates per unit physical time. With the latter determining the selection pressure on mass, it follows that the ∂ ln *w/∂* ln *∊* relation of eqn 22 has a metabolic component

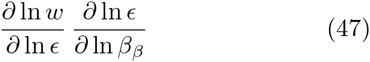

where pre-mass selection on metabolic pace is generating net energy for the selection of mass. Given the log-linear selection relation of eqn 23, and ∂ ln *∊/∂* ln *β_β_* = 1 from 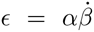, we find that primary selection on metabolic pace selects mass as a partial function of mass specific metabolism

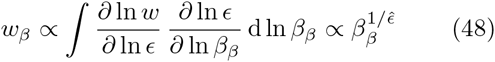

with the inverse

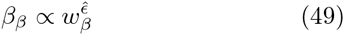

of the selection function being the component of the post-mass allometry for mass specific metabolism that evolves from pre-mass selection on metabolic pace.

Instead of dealing with mass as a joint parameter of several sub-components, like *w* = *w_α_w_β_*, I will express the evolutionary dependence of mass on *α* and *β_β_*, and the more usual inverse allometric correlations, as functions of total mass *w*. For this, let the dependence of *w* on *β_β_* that is captured by the allometry of eqns 48 and 49, be expressed by the following allometry

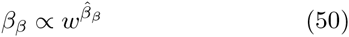

Note that the relative dependence of total mass on mass specific metabolism, as described e.g. by the *w_β_/w-*ratio, is expressed differently by eqns 49 and 50. In eqn 49 we have an invariant exponent 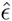, with the dependence of mass on mass specific metabolism being captured by the *w_β_* component that is directly dependent on *β_β_*. In eqn 50 it is instead the 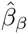 exponent that will change as a function of the *w_β_/w* ratio (eqn 29), to make any given dependence of total mass *w* on *β_β_* consistent across the range of possible *β_β_* values.

Now, if we insert eqn 50 into eqn 46, we find that the metabolic-rescaling components of the life history traits can be expressed by allometric relations of total mass

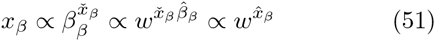

with 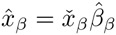.

#### 3.2.1 Allometric deduction

To deduce the metabolic-rescaling exponents, from the inverse relationship between pace and biotic time periods, we have

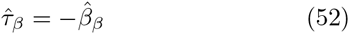

And from 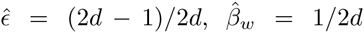 and the *∊* ∝ *β_β_ β_w_* constraint of eqn 28 that links net energy, resource handling, and pace we have

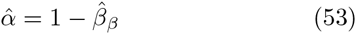

From the invariant selection optimum of density regulation *h^**^* ∝ (*v/β*)*^d^* (eqn 18), and a foraging speed 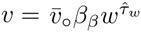 that is defined by the mass-rescaling for lifespan 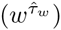 and the metabolism rescaling for metabolism *β_β_*, eqn 15) we have 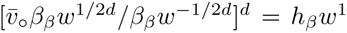 given 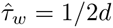, and 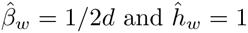. When solved for the mass-rescaling intercept for the home range, we find that it is an invariant intercept

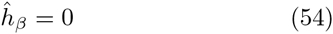

that maintains the population at the selection optimum.

From the 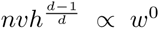 invariance of interference competition (eqn 19), with 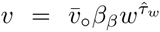 we have 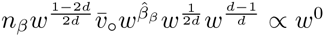 given 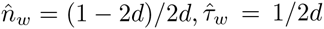 and 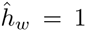. When solved for the mass-rescaling intercept of abundance, we find that it is a population density that scales inversely with the metabolic intercept

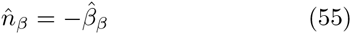

that maintains a body mass invariant level of interference competition in the population. This rescaling will also maintain an invariant exploitation of the resource (eqn 13), with an invariant use of energy by the population.

If we turn to the population dynamic equilibrium 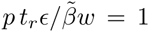 it implies 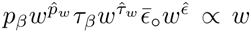, given 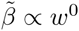. With 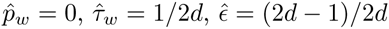 and 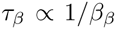 we find that it is a survival mass-rescaling intercept that is proportional to the metabolic intercept

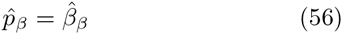

**Table 2:** Theoretical allometries. Allometric exponents 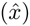 as they evolve from allometric rescaling given primary selection on metabolism and mass. The exponents depend on the dimensionality of the interactive behaviour (1D, 2D or 3D), and on the 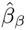 exponent that describes the relative importance of mass specific metabolism for the net energy of the organism.

| a) One dimensional interactions ($\hat{\epsilon} = 1/2$) | | | | | | | | |
| --- | --- | --- | --- | --- | --- | --- | --- | --- |
| $\hat{\beta}_\beta$ | $\hat{\alpha}$ | $\hat{\beta}$ | $\hat{\tau}$ | $\hat{p}$ | $\hat{R}$ | $\hat{r}$ | $\hat{h}$ | $\hat{n}$ |
| 0 | 1 | $-\frac{1}{2}$ | $\frac{1}{2}$ | 0 | 0 | $-\frac{1}{2}$ | 1 | $-\frac{1}{2}$ |
| $\frac{1}{2}$ | $\frac{1}{2}$ | 0 | 0 | $\frac{1}{2}$ | $-\frac{1}{2}$ | 0 | 1 | -1 |
| 1 | 0 | $\frac{1}{2}$ | $-\frac{1}{2}$ | 1 | -1 | $\frac{1}{2}$ | 1 | $-\frac{3}{2}$ |

| b) Two dimensional interactions ($\hat{\epsilon} = 3/4$) | | | | | | | | |
| --- | --- | --- | --- | --- | --- | --- | --- | --- |
| $\hat{\beta}_\beta$ | $\hat{\alpha}$ | $\hat{\beta}$ | $\hat{\tau}$ | $\hat{p}$ | $\hat{R}$ | $\hat{r}$ | $\hat{h}$ | $\hat{n}$ |
| 0 | 1 | $-\frac{1}{4}$ | $\frac{1}{4}$ | 0 | 0 | $-\frac{1}{4}$ | 1 | $-\frac{3}{4}$ |
| $\frac{1}{4}$ | $\frac{3}{4}$ | 0 | 0 | $\frac{1}{4}$ | $-\frac{1}{4}$ | 0 | 1 | -1 |
| $\frac{1}{2}$ | $\frac{1}{2}$ | $\frac{1}{4}$ | $-\frac{1}{4}$ | $\frac{1}{2}$ | $-\frac{1}{2}$ | $\frac{1}{4}$ | 1 | $-\frac{5}{4}$ |
| 1 | 0 | $\frac{3}{4}$ | $-\frac{3}{4}$ | 1 | 1 | $\frac{3}{4}$ | 1 | $-\frac{7}{4}$ |

Table 2: **Theoretical allometries.**
| c) Three dimensional interactions ($\hat{\epsilon} = 5/6$) | | | | | | | | |
| --- | --- | --- | --- | --- | --- | --- | --- | --- |
| $\hat{\beta}_\beta$ | $\hat{\alpha}$ | $\hat{\beta}$ | $\hat{\tau}$ | $\hat{p}$ | $\hat{R}$ | $\hat{r}$ | $\hat{h}$ | $\hat{n}$ |
| 0 | 1 | $-\frac{1}{6}$ | $\frac{1}{6}$ | 0 | 0 | $-\frac{1}{6}$ | 1 | $-\frac{5}{6}$ |
| $\frac{1}{6}$ | $\frac{5}{6}$ | 0 | 0 | $\frac{1}{6}$ | $-\frac{1}{6}$ | 0 | 1 | -1 |
| $\frac{1}{2}$ | $\frac{1}{2}$ | $\frac{1}{3}$ | $-\frac{1}{3}$ | $\frac{1}{2}$ | $-\frac{1}{2}$ | $\frac{1}{3}$ | 1 | $-\frac{4}{3}$ |
| 1 | 0 | $\frac{5}{6}$ | $-\frac{5}{6}$ | 1 | -1 | $\frac{5}{6}$ | 1 | $-\frac{11}{6}$ |

that maintains the balance of the population dynamic equilibrium. This increased probability of surviving to reproduce with a pre-mass increase in mass specific metabolism reflects a decline in the mortality rate in biotic time; a decline that may reflect a shortening of the physical time period where the individual is exposed to extrinsic mortality factors.

With *R^*^* = 1*/p^*^* it follows that the mass-rescaling intercept of lifetime reproduction is inversely related to the intercept of the metabolic rate per unit body mass 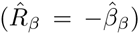, and from *r* = ln(*pR*)*/t_g_* we have 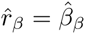. These changes in the mass-rescaling intercepts from metabolic-rescaling are listed in Table 1b.

### 3.3 Final allometries

Given the deduced exponents in Table 1 for the allometric rescaling with mass and metabolism we can calculate the 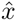 exponents of the final post-mass scaling of the life history with mass for a variety of situations. I list these exponents in Table 2 for interactive behaviour in one, two and three spatial dimensions.

Apart from the home range exponent that is always one, the exponents depend on the 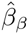 exponent that describes the relative importance of metabolic pace for net resource assimilation and, thus, also for the selection of mass. This implies a post mass exponent for mass specific metabolism 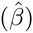 that increases with the relative importance of metabolism for the evolution of mass. This is illustrated in Table 2, where 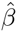 is 3/4 for 2D, 5/6 for 3D, interactions in the extreme cases where all of the variation in *∊*, and thus also body mass, is caused by variation in the pre-mass component of mass specific metabolism 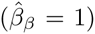. At the other extreme where all the body mass variation is caused by variation in resource handling and/or resource availability 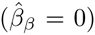, the metabolic exponent takes the more well-known values of −1/4 (2D) and −1/6 (3D). For an intermediate case with a similar importance of handling and pace 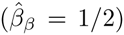 we have 1/4 for 2D and 1/3 for 3D. The cases where mass specific metabolism is independent of mass 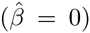 is also shown. The latter depends on the spatial dimensionality of the interactive behaviour, with 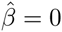 for 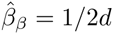.

The majority of post-mass exponents are given as fractions, where 2*d* is the common denominator, with the most well-known set of allometric exponents for large bodied species, i.e., the set with 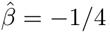, evolving for two dimensional interactions when all the variation in body mass is caused by variation in resource handling/availability. This case has net energy that scales to the 3/4 power of mass, biotic periods that scales to the 1/4 power, population densities that scale to the −3/4 power, and population growth that scales to the −1/4 power of mass. The corresponding exponents for one dimensional interactions are ±1/2, while the ±1/4, and ±3/4, exponents are exchanged with ±1/6, and ±5/6, exponents for interactive behaviour in three spatial dimensions.

## 4 Allometric evidence

In the evaluation of evidence in favour of my model I will focus on the average and commonly observed empirical exponents across the tree of life. With an underlying mechanism that is based on ecological inter actions, we can expect deviations from the predicted exponents in some phylogenetic lineages and competitive guilds. The model is however not currently developed to explain these deviations, and I will therefore not consider deviating exponents in detail. The point to make here is not that all empirical exponents resemble the predicted; because this is expected not to be the case. The point is instead whether the predicted exponents and their transitions are commonly observed and in general agreement with empirical data.

### 4.1 Prokaryotes

DeLong et al. (2010) estimated an allometric exponent for mass specific metabolism of 0.96±0.18 across active prokaryotes, and 0.72 ± 0.07 across inactive. Given the theoretical allometries in Table 2, this indicates three dimensional ecology with an average exponent of 0.84, and body mass variation that evolves from primary variation in mass specific metabolism.

### 4.2 Protozoa

Based on the data from Makarieva et al. 2008, DeLong et al. (2010) estimated the average exponent across the complete range of inactive protozoa to be −0.03 (±0.05), and that of active protists to be 0.06 (±0.07). These exponents are smaller than the exponent for prokaryotes and larger than the typical −1/4 and −1/6 exponents for multicellular animals.

By excluding the four smallest protozoa with exceptionally high metabolic rates from the data of Makarieva et al. (2008), and by least-squares fitting a third degree polynomial to the remaining data for inactive protozoa (*n* = 48), I obtained point estimates of the body mass exponent for mass specific metabolism that declined from 0.61 across the smallest [log *w*(kg) = −13.5], over zero across intermediate [log *w*(kg) = −11], to a minimum of −0.20 among the largest protozoa [log *w*(kg) = −8.0].

### 4.3 Multicellular animals

The often reported −1/4 exponent for mass specific metabolism in multicellular animals indicates that a major component of the body mass in this group is selected from primary variation in the handling and/or density of the underlying energetic resources.

#### 4.3.1 Life history and ecological traits

For any group of organisms, it is the terrestrial vertebrates that have been subjected to most allometric studies, and in Table 3 I show the predicted 2D exponents for the case of Kleiber scaling, together with some of the commonly observed exponents in mammals, reptiles and birds. It can be concluded that a reasonable resemblance exists between the theoretical and empirical exponents across traits ranging from metabolism, lifespan, survival and reproduction over population growth to ecological traits like the home range of individuals and the densities of populations.

**Table 3:** Empirical exponents for different traits. The theoretical 2D exponents compared with empirical exponents for reptiles, birds and mammals. From Witting (1997), except 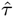 for reptiles from Calder (1984).

| $\hat{x}$ | 2D | Reptiles | Birds | Mammals |
| --- | --- | --- | --- | --- |
| $\hat{\beta}$ | -0.25 | -0.24 | -0.26 | -0.26 |
| $\hat{\tau}$ | 0.25 | 0.23 | 0.18 | 0.25 |
| $\hat{p}$ | 0.00 | - | 0.01 | - |
| $\hat{R}$ | 0.00 | - | 0.00 | -0.03 |
| $\hat{h}$ | 1.00 | 0.95 | 1.16 | 0.99 |
| $\hat{n}$ | -0.75 | -0.77 | -0.75 | -0.78 |

### 4.3.2 2D versus 3D

Another essential prediction is the transitions in the allometric exponents across species that differ in the spatial dimensionality of their interactive behaviour.

To examine the evidence for these transitions, I have in Table 4a listed allometric exponents for mass specific metabolism 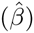 across the species of major taxonomic groups, with the empirical 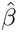 estimates being used to classify the taxa as having behavioural interactions in 2D or 3D (assuming that 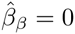). Most of these exponents are either the average exponent, or the exponent based on the largest sample size, from the tables in Peters (1983). I found no convincing 1D cases, but the exponents for mass specific metabolism are in general agreement with the hypothesis that they evolve from the underlying spatial dimensionality of intra-specific interactions.

The interactions of most terrestrial and benthic taxa are classified as 2D, and those of pelagic and tree living taxa as 3D. This overall separation is likely to reflect that most terrestrial and benthic animals are constrained to behavioural interaction in the two horizontal dimensions, while pelagic and tree living species have an extra vertical dimension in which to forage and interact.

With the available data the differentiation in the allometric exponents is maybe clearest in mammals, that are dominated by ground living 2D species and an overall exponent for field mass specific metabolism of −0.25 ± 0.03 (Savage et al. 2004). Yet pelagic taxa like Cetacea, Pinnipedia and Sirenia have a 3D exponent of 0.16 ± 0.02 for lifespan (Witting 1995), and primates, where the majority of species are tree-living, are also classified as 3D with an exponent of −0.19 ± 0.03 for mass specific metabolism (Genoud 2002).

**Table 4:** Empirical 2D and 3D allometries. a): Taxa classified as 2D or 3D from estimates of the 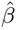 exponent (±SD. **b**): Exponents for 2D - 3D transitions: 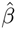, as defined by the average exponents of Table a; 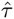, terrestrial versus pelagic mammals (Nowak, 1991); 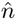, 2D-3D classification and exponents from Pawar et al. (2012). a:Peters (1983) [1:estimate with largest sample size; 2:average estimate; 3:average estimate, but −1 outlier]; 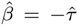, Nowak (1991); c:Genoud (2002).

| a) $\hat{\beta}$ exponent classification | | | |
| --- | --- | --- | --- |
| Group | 2D | 3D |  |
| Mammals (all)<sup>a,1</sup> | $-0.26 \pm 0.01$ | | |
| Pelagic mammals<sup>b</sup> | | $-0.16 \pm 0.02$ | |
| Primates<sup>c</sup> | | $-0.19 \pm 0.03$ | |
| Bats<sup>a</sup> | $-0.26$ | | |
| Birds<sup>a,1</sup> | $-0.26 \pm 0.01$ | | |
| Reptiles<sup>a,2</sup> | $-0.24 \pm 0.02$ | | |
| Snakes<sup>a,3</sup> | $-0.26 \pm 0.04$ | | |
| Lizards<sup>a,2</sup> | | $-0.18 \pm 0.02$ | |
| Turtles<sup>a,2</sup> | | $-0.14 \pm 0.03$ | |
| Frogs<sup>a,2</sup> | $-0.29$ | | |
| Salamanders<sup>a,1</sup> | | $-0.18 \pm 0.02$ | |
| Fishes<sup>a,3</sup> | | $-0.20 \pm 0.01$ | |
| Molluscs<sup>a</sup> | $-0.25$ | | |
| Jellyfishes<sup>a</sup> | | $-0.15$ | |

Table 4: **Empirical 2D and 3D allometries.**
| b) 2D - 3D transitions |  |  |  |
| --- | --- | --- | --- |
| d | $\hat{\beta}$ | $\hat{\tau}$ | $\hat{n}$ |
| 2D | $-0.26 \pm 0.02$ | $0.25 \pm 0.04$ | $-0.79 \pm 0.09$ |
| 3D | $-0.17 \pm 0.02$ | $0.16 \pm 0.02$ | $-0.86 \pm 0.08$ |

Other plausible 2D-3D classifications of interactions from the exponent of mass specific metabolism include reptiles, snakes, frogs and mollusks as 2D, and fishes, jellyfishes and salamanders as 3D (Table 4a). However, there are also a few less evident cases. Why, e.g., do birds and bats group as 2D? Although birds and bats move freely in 3D, it is most likely the packing of their breeding and/or feeding territories that are essential for the evolution of the allometric exponents, and this packing is 2D in many, if not most, cases. A splitting of birds and bats into those that compete for territories and food in 2D and 3D would be desirable, but beyond this study that is based on empirical literature estimates of allometric exponents.

The dimensionality estimates of the intra-specific interactions from metabolic exponents in lizards, snakes, reptiles and turtles might also benefit from a split into 2D (e.g., ground) and 3D (e.g., tree or pelagic) living species, with the reported exponents reflecting 3D for lizards and turtles, and 2D for reptiles and snakes.

The split between 2D and 3D interactions has been observed not only in the exponent for mass specific metabolism, but also in exponents for lifespan and population density (Table 4b), and it seems to relate also to allometries of community ecology (Pawar et al. 2012).

### 4.3.3 Invariant interference

Interaction levels have hardly been reported for any species, but evidence for or against the existence of invariant interference competition among large bodied species may be examined by a comparison of empirical allometries. From eqn 16 we expect a level of interference that is proportional to 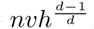, with 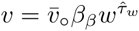 (eqn 15). Concentrating on two dimensional foraging with 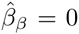 we find a level of interactive competition that scales as

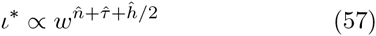

Across 2D species in major taxonomic groups 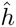 is usually approximately 1, 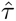 approximately 1/4, and 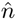 around −3/4 (Table 4c; Peters 1983; Calder 1984; Damuth 1987; Nee et al. 1991; Witting 1995). From the exponents in Table 4c we find that 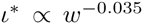 for mammals, 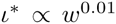 for birds, and 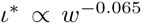 for reptiles. The theoretically deduced invariance with respect to body mass is not contradicted by data.

#### 4.3.4 Mass-rescaling intercepts

Evidence on the scaling of the intercepts of the traditional mass-rescaling allometries with Kleiber scaling is not as clear as for the mass-rescaling exponents. But with ectotherms having field metabolic rates that are 12 to 20 times smaller than in similar sized endotherms (Nagy 2005), from Table 1b we expect ectotherms to have longer lifespans and to be more abundant than similar sized endotherms, and this is generally the case (Peters 1983; Currie and Fritz 1993; deMagalhães et al. 2007). The predicted inverse relationship between lifetime reproduction and the mass-rescaling intercept for mass specific metabolism is also in agreement with fecundity estimates that are about ten times higher in reptiles than in mammals (Peters 1983) reflecting, as predicted, a higher probability to survive to reproduce in mammals.

## 4 Discussion

In this paper I added primary selection on metabolism to the ecological selection model of Witting (1995), and showed that the joint selection of metabolism and mass can explain a wide range of allometric exponents that are observed across the three of life.

### 5.1 Mass specific metabolism

For this I separated the resource assimilation parameter of the original model into resource handling and the pace of handling. This pace generates gross energy, and by defining net energy as the difference between gross energy and the total metabolism of the organism, I found the mass specific work of handling to be selected as mass specific metabolism. This implies primary selection for an increase in mass specific metabolism, and this generates the net energy that is a pre-condition for the selection of mass and body mass allometries.

Given the importance of metabolism for animals (e.g., Lotka 1922; Pearl 1928; Maynard Smith and Szathmáry 1995; Brown et al. 2004) it is somewhat surprising that the metabolic rate has been the forgotten life history character, with not even a single model on its selection in seven review books on life history evolution (Charlesworth 1980, 1994; Roff 1992, 2002; Stearns 1992; Bulmer 1994; Stearns and Hoekstra 2000; but see Witting 2003; Barve et al. 2014; Artacho and Nespolo 2009; Boratyñski et al. 2010; Versteegh et al. 2012).

The evolution of metabolism has instead been studied as a molecular process in relation to the origin of life (Horowitz 1945; Miller 1953; Haldane 1954; Oparin and Clark 1959; Ponnamperuma and Chela-Flores 1993; Chela-Flores et al. 1995; Baltscheffsky 1996; Cunchillos and Lecointre 2003; Ferry and House 2006; Melendez-Hevia et al. 2008; Fernando and Rowe 2007, 2008; Fry 2011; Marakushev and Belonogova 2013). And it has been examined empirically, where metabolism in natural species is linked to a range of extrinsic factors like temperature (e.g., McNab and Morrison 1963 Lovegrove 2003; Wikelski et al. 2003; Careau et al. 2007; Jetz et al. 2007; White et al. 2007), primary production (Mueller and Diamond 2001; Bozinovic et al. 2007, 2009), rainfall (Lovegrove 2003; Withers et al. 2006; White et al. 2007), and diet (McNab 2003; Anderson and Jetz 2005; Muñoz-Garcia and Williams 2005). Fitness components like survival and reproduction have also been found to correlate with metabolism in a wide range of species (see e.g. Table 4 in White and Kearney 2013), and these correlations tend to indicate a stabilizing selection (Artacho and Nespolo 2009; White and Kearney 2013). This suggests that natural species are optimized towards some central metabolic value that is given by the current evolutionary state of the species in a given environment. Yet, the study of empirical fitness correlations with metabolism is insufficient in itself as it does not reveal the underlying selection of the attracting metabolic rate.

Another widespread view is that of metabolic ecology (Brown et al. 2004; Sibly et al. 2012; Humphries and McCann 2014; Padfield et al. 2016), where metabolism is seen to determine the rate at which the organism assimilates, transforms and expends energy. Instead of formulating this as a life history trait that is selected by natural selection, metabolic ecology treats metabolism as a passive physical parameter (Glazier 2015) that is determined by temperature and the geometry and physics of resource transportation networks (West et al. 1997, 1999; Gillooly et al. 2001). While metabolism is indeed essential for the pace of physiological and ecological processes, and while physical factors may constrain the metabolic rate, metabolic ecology is insufficient in itself because there are no physical laws that will explain the elevated rates of metabolism in mobile organisms in general, and in birds and mammals in particular.

In this paper, I showed theoretically that mass specific metabolism is likely to be selected as the life history character that determines the pace of the net resource assimilation that generates net energy for self-replication and the selection of mass. This provides an overall direction where unconstrained selection is generating an exponential increase in mass specific metabolism and the net energy of the organism. Associated with this increase there is a metabolic rescaling of the rate dependent life history characters, and this rescaling is affecting the body mass allometries because the selected mass is dependent on the energy that is generated from the selected increase in mass specific metabolism.

### 5.2 Mass-rescaling selection

The well-known 1/4 exponents of Kleiber scaling, however, was found to be selected by a mass-rescaling response to the evolutionary changes in mass. While implicit in my original allometric model (Witting 1995), the first formal description of mass-rescaling selection is presented in the current paper. This selection emerges from the energetic trade-off between mass, metabolism and the time that it takes to grow a full-sized offspring, and it selects for a secondary decline in mass specific metabolism as a response to an increasing mass.

With an increasing mass, reproduction will decline as it takes longer to grow larger offspring when the rate of mass specific metabolism is the same. But the reproductive rate can be maintained if metabolism is reduced so that more of the parental energy is allocated into the mass of the offspring. This selection, however, goes not only contrary to the pre-mass selection for increased pace, but the selection is also dependent upon the organism energy that is generated by the pre-mass selection and, thus, it cannot just reduce the metabolic pace as this would eliminate the net energy that generates the selection. The evolutionary solution is an inverse scaling between mass specific metabolism and biotic time periods, as this will maintain organism energy, reproduction and metabolic pace constant on the per generation time-scale of natural selection, while the three traits are declining in physical time with an evolutionary increase in mass.

While it is the energetic trade-off between mass, metabolism and time that makes metabolism and biotic time periods respond to the evolutionary changes in mass, it is primarily the ecological geometry of optimal density regulation that was found to determine the actual values of the response as defined by allometric exponents. The mass induced changes in metabolic pace and biotic time is affecting the foraging process, and consequently the selection for a home range that satisfies the conditions of optimal density regulation. The dependence of interactive competition and local resource exploitation on the spatial dimensionality (*d*) of organism behaviour is then transferred by the selected density regulation optimum to the values of the allometric exponents, with the 1/4 exponent being the 2D case of the more general 1/2*d*.

### 5.3 Diverse allometries

While mass-rescaling selection was found to be responsible for the evolution of the often observed Kleiber scaling, it was also found that a broader understanding of allometries is dependent on the inclusion on primary selection of metabolic pace.

The −1/4 exponent of Kleiber scaling was found to be restricted mainly to the taxa of multicellular animals that evolve masses from intra-specific interactions in two spatial dimensions, when these taxa diversity by species that evolve into a multitude of ecological niches. Such a diversification will allow the pre-mass variation in resource handling and resource availability to dominate pre-mass variation in metabolic pace, with the result that post-mass allometries evolve primarily from mass-rescaling with a −1/4 exponent. The corresponding 3D exponent is −1/6, and the −1/4 *↔ −*1/6 transition is observed quite commonly between terrestrial and pelagic taxa (Table 4a).

A body mass invariance of mass specific metabolism across major taxa from prokaryotes to mammals (Makarieva et al. 2005, 2008; Kiørboe and Hirst 2014) was instead explained by a macro evolution where natural selection has taken mass specific metabolism to an upper bound of the evolved metabolic pathways.

An average exponent around 0.84 for mass specific metabolism across the masses of prokaryotes (DeLong et al. 2010) is consistent with 3D selection and body mass variation that is selected primarily from variation in the pre-mass component of mass specific metabolism.

And an observed decline in the exponent for mass specific metabolism from 0.61 over zero to −0.20 in protozoa (Section 4.2) is predicted by a gradual change in the natural selection of mass; suggesting that the mass of the smallest protozoa is selected from primary variation in mass specific metabolism, while the mass of the largest is selected from primary variation in the handling and/or density of the underlying resource.

These results indicate that unicellular protozoa may evolve as a continuum that spring from the selection mechanism in prokaryotes (where mass is selected from primary variation in mass specific metabolism), and undergoes a gradual change with an increase in mass towards the selection mechanism in multicellular animals (where mass is selected from primary variation in the handling and/or density the underlying resource). This apparent shift in the natural selection of mass across the tree of life is studied by Witting (2016a), who shows that lifeforms from virus over prokaryotes and larger unicells to multicellular animals follow as a unidirectional unfolding of the allometric model that I have proposed here.

Inter-specific exponents that are observed across natural species may nevertheless differ from the predictions. As the predicted exponents evolve from the spatial dependence of density regulation on traits like metabolism, foraging speed, home range, population density and mass, it is only natural to expect some variation in the exponents. While the base-case description in Section 2.3 of this spatial density dependence appears to be representative for the average exponents across a broad range of mobile organisms, we would expect some variation in the density dependent ecology and this may result in deviating allometric exponents.

Reasons for exponents that deviate from the predictions in this paper may include inter-specific interactions that bias the resource distribution with mass in local communities (Brown and Maurer 1986; Nee et al. 1991) away from the invariance that applies at scales where the competitive effects from inter-specific interactions are minimal. Inter-specific interactions may also alter the optimum of the density dependent foraging behaviour, with possible deviations in the allometric exponents. Small islands, let it be real islands or habitat islands on a mainland, may limit the size of home ranges causing deviations in allometric exponents. Finally, the deduction in this paper assumes some invariance in the life history, and is therefore not strictly valid across gradients where, e.g., the age of maturity is correlated with mass when measured in biotic time. Empirical examples of inter-specific exponents that deviate from the expected can be found in the tables of Peters (1983), and they are discussed by McNab (1988, 2008), Lovegrove (2000), Glazier (2010, 2015), Kolokotrones et al. (2010) and others.

### 5.4 Parsimonious evolution

The proposed selection was elaborated from the foraging ecology in one of the earliest deductions of allometric exponents (Witting 1995), and it explains allometric exponents from primary selection on metabolism and mass. Yet, the more widespread view has seen the metabolic exponent as a mechanical or evolutionary consequence of physiological constraints. Let it be from a surface rule in four spatial dimensions (Blum 1977), resource uptake and use at cell or body surfaces (Davison 1955; Patterson 1992; Makarieva et al. 2003), tissue demands for resources (McMahon 1973; Darveau et al. 2002), resource demand with cellular and demographic constraints (Kozlowski and Weiner 1997; Kozlowski et al. 2003a,b), resource demand and exchange (Sibly and Calow 1986; Kooijman 2000; Banavar et al. 2002a,b), geometric constraints on resource transportation systems (West et al. 1997, 1999a,b; Banavar et al. 1999; Dodds et al. 2001; Dreyer and Puzio 2001; Rau 2002; Santillán 2003), thermodynamic constraints at the molecular level (Fujiwara 2003), quantum mechanical constraints on proton and electron flow in metabolic pathways (Demetrius 2003, 2006), ecological metabolism constrained by physical limits on metabolic fluxes across surface-areas in relation to volume dependent resource demands (Glazier 2005, 2010), or from scaling in the four dimensions of space and time (Ginzburg and Damuth 2008).

As none of these alternative models explain how natural selection generates the evolution of the required co-existence of mass and metabolism, they fail to explain the span of organisms that are a pre-condition for the existence of allometries. Several of the studies have though identified empirical 1/4 exponents for diverse physiological processes, and this may illustrate just how deep into the physiology the ecological constraints of optimal density regulation is selected. Resource transportation networks, e.g., are selected to supply the organism with energy. One potential solution is a fractal network of branching tubes (West et al. 1997) that can easily be adjusted to comply with a −1/4 exponent that is selected by the ecological geometry of foraging (Witting 1998).

## Appendix

### A Life history

The conservation of energy is constraining the possible trait space of the life history (e.g., Charlesworth 1980; Roff 1992, 2002; Stearns 1992; Bulmer 1994; Stearns and Hoekstra 2000), with some of the more essential constraints and trade-offs being described in this section.

#### A.1 Biological time

To include time in this conservation, let the age of maturity (*t_m_*, age of first reproductive event in physical time) divide the potential lifespan (*t_l_*) into a juvenile period (*t_j_* from age 0 to *t_m_*) and a reproductive period (*t_r_* from *t_m_* to *t_l_*). The generation time may then be seen as the average age of reproduction 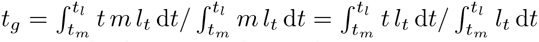, given a constant reproductive rate *m* over the reproductive period, and the probability 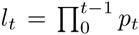 that an individual will survive to age *t*, with *p_t_* being the probability to survive from age *t* to age *t* + 1.

As mass specific metabolism is selected as the pace of the biochemical, physiological and ecological processes of resource assimilation, we can follow Pearl (1928) and others like Brody (1945), Hill (1950), Stahl (1962) and Calder (1984), and consider biological time as inversely related to mass specific metabolism,

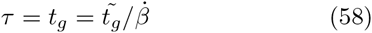

with *τ* being the per generation time-scale of natural selection, and 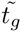 an invariant scaling between generation time and metabolic pace 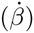. The relation between a period *x*, or age, of an organism in physical (*t*) and biotic time (*τ*) is thus

**Table 5:**
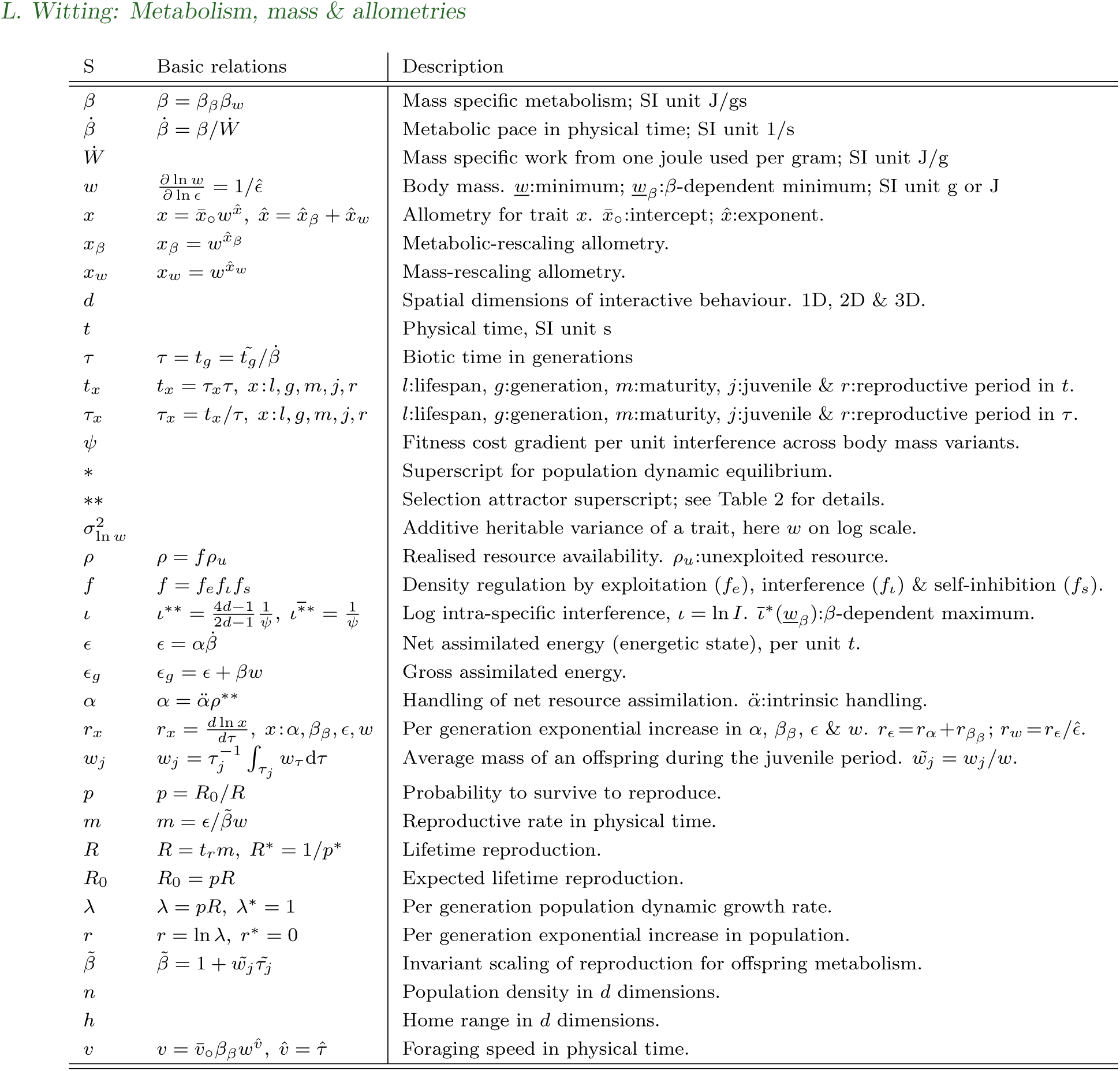
Symbols (S) with basic relationships.

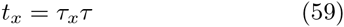

with *τ_g_* = *t_g_*/*τ* = 1.

#### A.2 Energy, metabolism and mass

To formulate the energetic trade-off between metabolism and mass, let the body mass (*w*) be defined by all the biochemical energy that it takes to build the mass. That is, let it be the biochemical energy of the matter that forms the mass, plus the energy that is used by anabolic metabolism to build the organism from smaller molecules.

Given complete parental investment, we have that the average mass of an offspring (*w_j_*) during the juvenile period (*t_j_*) where it is reared by the parents is

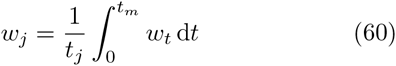

The total energetic investment in each offspring is then the final adult mass *w* of the offspring at the time of independence, plus the energy *w_j_t_j_* that is metabolized by the offspring during the juvenile period

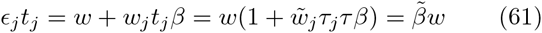

where *∊_j_* is the energetic investment in the offspring per unit time, and

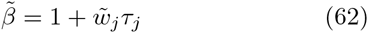

is invariant for invariant *τ_j_* and 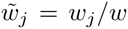. The total investment in each offspring is then reducing to a quantity that is proportional to mass and independent of mass specific metabolism.

The anabolic energy that is used to build the offspring is supplied by the net energy (*∊*) of the parent, and it is defined here to be part of the mass (*w*) of the offspring, instead of being part of the metabolism (*β*) of the offspring.

#### A.3 Reproduction

The energetic constraint on reproduction is described by a quality-quantity trade-off where the parent is constrained to produce many small, or a few large, off-spring. With *∊* being the energy available for reproduction, the reproductive rate on the time-scale of the juvenile period is *m_t__r_* = *∊/∊_j_*. Yet, with the total energetic investment in each offspring being defined by eqn 61, the reproductive rate in physical time

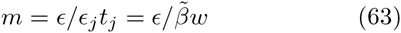

is proportional to the net energy and inversely proportional with mass, with total lifetime reproduction

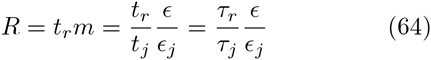

being constrained by the *τ_r_τ_j_* and *∊/∊_j_* ratios.

#### A.4 Survival

As the expected lifetime reproduction of a new-born is

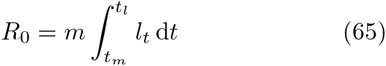

we find that the probability that a new-born individual will survive the complete reproductive period is

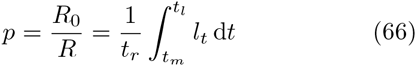

For this relation we note that when the *l_t_* function is invariant in biotic time, i.e., when *l_t/t__r_* is invariant, it follows that the 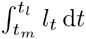 integral reduces to *c_l_t_r_* where *c_l_*, and thus also *p*, are invariant. This is our base-case expectation if survival is determined entirely by intrinsic processes. On top of this there is extrinsic mortality, and if this is about constant in physical time we might expect *p* to be more or less inversely related to *t_r_* with more individuals dying before they reach a given age in biotic time.

### B density regulation

Some of the most essential constraints for the evolution of the allometric exponents are imposed by the ecological geometry of the density dependent foraging and interactive competition that is regulating the dynamics of the population. This was first described by Witting (1995), and his model is given here in a more general version.

The potential importance of density dependence for life history evolution has been realized since MacArthur (1962); first with the verbal formulation of *r* and *K* selection (Pianka 1970; Stearns 1976, 1977; Parry 1981), and then with its more formal mathematical theory (Anderson 1971; Roughgarden 1971; Charlesworth 1971; Clarke 1972), with frequency independent selection for an increase in *r* or *K*, as noted by Fisher (1930) when he formulated his fundamental theorem of natural selection (Witting 2000a, 2002b).

It was found later that the incorporation of the frequency dependent interactive behaviour is essential for a general understanding of density dependent selection (e.g. Abrams and Matsuda 1994; Mylius and Diekmann 1995; Metz et al. 1996; Witting 1997; Heino et al. 1998; Gyllenberg and Parvinen 2001; Dercole et al. 2002; Dieckmann and Metz 2006).

#### B.1 The overall regulation

To formulate this regulation, let net energy 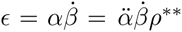 be defined for the realized resource density (*ρ*) in a population that has evolved to the relevant selection attractor for mass (e.g., *\*\**). The density regulation of net resource assimilation may thus be given by *∊ρ/ρ^**^*, with *ρ* being the density (*n*) regulated *f*(*n*) resource

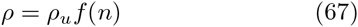

that declines monotonically from *ρ_u_* to zero as *n* increases from zero to infinity, with *ρ_u_* being *ρ* at the lower limit *n* ≈ 0, assuming no depensation. The equilibrium abundance

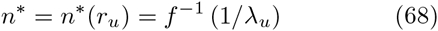

is constrained by the 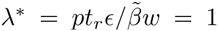 relation of eqn 11 where *∊* = *∊_u_f*(*n*), and it increases monotonically with the maximal growth rate 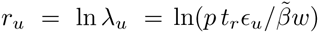, with 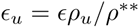.

As an approximation of density regulation in the surroundings of the equilibrium I assume log-linearity

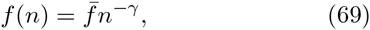

where 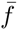 is a constant, and

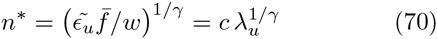

the equilibrium with 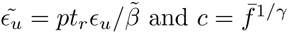.

Let the regulation function *f* be a joint function of three sub-components

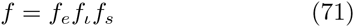

where *f_e_* is regulation by exploitative competition, *f_l_* is regulation by interference competition, and *f_s_* is local resource exploitation by self-inhibition. Self-inhibition is not density regulation in itself, but it is needed for the evolution of a realistic density regulation (Section B.5).

#### B.2 Exploitative competition

Density regulation by exploitative competition occurs through the reduction in resource density caused by the consumption of resources by the population. And with the energy that is used by the juvenile component *n_j_* being supplied by the parental generation (*n_a_* = *n − n_j_*), it is a function of the gross resource consumption (eqn 2) of the adult component

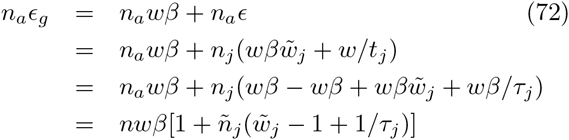

where the net energy of the adult component (*n_a_∊*) is used by the juvenile component (eqn 61) for metabolism 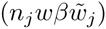 and mass (*n_j_w/t_j_*), with 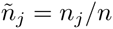.

As it is reasonable to assume that 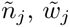 and *τ_j_* are invariant of mass, we obtain

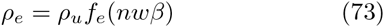

as a general expression of the exploitation function, with *f_e_* declining monotonically from one to zero as *nwβ* increases from zero to infinity, and *ρ_e_* being the density of the exploited resource. The resource density *ρ_e_* is different from the realized resource *ρ* = *ρ_u_f* = *ρ_e_f_l_f_s_*, as the latter refers to the resource component that can effectively be exploited after the cost of interference and local exploitation.

As an approximation in the surroundings of *n^*^* I use

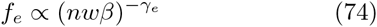

with *γ_e_* being the density regulation parameter of exploitative competition.

#### B.3 Interactive competition

Density regulation by interactive competition reflects the costs of interference, with the cost to an individual including both the cost of losing access to resources that are controlled by competitively superior individuals, and the cost of using time and energy on competitive interactions with other individuals.

Where it is the bias in the cost of interference across the individuals in the population that is essential for interactive selection, it is the average cost that is essential for density regulation. This regulation is broken up into two components, the density dependence function

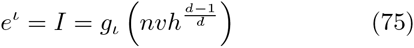

that determines the level of interactive competition, on ordinary (*I*) and log (*ι* = ln *I*) scale, from, among others, the density of the population, and the regulation function

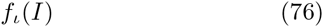

that declines monotonically from one to zero as *I* increases from zero to infinity.

To identify the 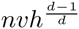 component of eqn 75 we note that the frequency of competitive encounters per individual is expected to be proportional to the degree of overlap between the home ranges of the individuals in the population, times the frequency by which the individuals reuse the foraging tracks within the home range. It is this frequency that defines how often the individuals will meet in overlapping areas.

Home range overlap may be defined as the average home range (*h*) divided by the per capita availability of space, which is proportional to 1*/n*; the inverse of population density. Hence, overlap is proportional to *nh*.

The time between reuse of foraging tracks is the length of the tracks divided by the foraging speed (*v*). Given *d*-dimensional foraging, the length of the foraging tracks is expected to be proportional to the *d*th root of the home range, as empirically confirmed for mammals (Garland 1983; Calder 1984). Hence, the interval between track reuse scale as *h*^1*/d*^/*v*. Multiplying the frequency of track reuse (*v/h*^1/*d*^) with home range overlap (*nh*) we find the level of interference competition (eqn 75) to increase monotonically from zero towards infinity as 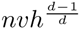 increases from zero to infinity.

The interference of eqn 75 applies when the interactive behaviour of the individuals in the population is unaffected by the resource density *ρ_e_*. While this may be expected for low energy organisms with a passive interactive behaviour, the level of interference in populations of high energy organisms might be inversely related to the underlying resource density 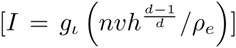 given that the individuals fight more often over resources when these are limited.

Foraging speed has been found to be proportional to lifespan and other organism periods on the body mass axis across species (Garland 1983; Calder 1984), with mechanistic explanations provided by Calder (1984). We expect also that foraging speed reflects metabolic pace, suggesting that the allometric intercept of foraging speed is proportional to the intercept of mass specific metabolism. Given these relations, we defined foraging speed as

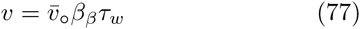

As an approximation in the surroundings of *n*^*^, I use

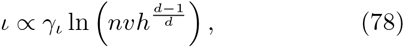

and

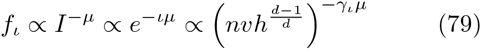

where *γ_ι_* is the density dependence of interference, and *μ* the average cost. Then, by inserting eqn 70 into eqn 78 we find the level of interference competition at the population dynamic equilibrium

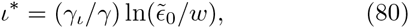

with 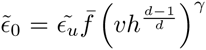. This level declines monotonically with *w*, with 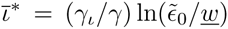 defining the maximum from an average minimum mass *<u>w</u>*.

#### B.4 Local resource exploitation

The resource assimilation of an individual is also influenced by the local resource exploitation of the individual itself. The availability of food along the foraging tracks of the individual is expected to be proportional to the time interval between the individuals reuse of foraging tracks, with longer intervals allowing more time for resource re-growth and/or dispersal into the area. Above we expressed this interval as *h*^1/d^*/v*.

But foraging self-inhibition by local resource exploitation is a relative term, relative to the frequency of re-harvesting when foraging tracks are infinitely long and never reused by the individual itself. In this case, re-harvesting is occurring because of the overall resource exploitation from all the individuals in the population, with a frequency that is expected proportional with biotic pace (*β*). When scaled accordingly we find self-inhibition expressed as

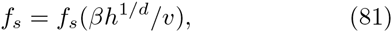

where *f_s_* is a downward bend function that increases monotonically from zero to one as the home range increases from zero to infinity.

As an approximation in the surroundings of an evolutionary equilibrium (Section B.5) I use

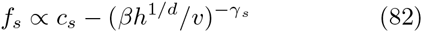

where *c_s_* is a scaling constant and *γ_s_* the strength of self-inhibition. And for the density regulation approximation 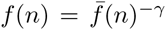, from eqns 74, 79, and 81 we have

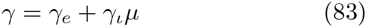

and

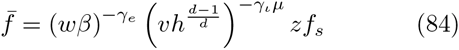

where *z* is a scaling parameter.

#### B.5 Selection of density regulation

Natural selection on the exploitative component of density regulation occurs through selection on the net assimilation of energy. This represents the ability of the organism to exploit resources, and it is covered by Section 2.3.

Relating to regulation by interactive competition 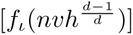 we note that, when this component is considered in isolation, we expect selection for home ranges that are so small that there are no interactions between individuals and no cost of interference. This does not coincide with natural conditions where interactions between individuals are common. Interactions, however, are expected because local resource exploitation [*f_s_*(*βh*^1/d^/*v*)] is counteracting interference, as it increases with a decline in home range. In fact, if we were only considering local resource exploitation, we would expect selection for infinitely large home ranges with no self-inhibition.

When the joint regulation 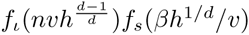 of the two functions are considered together, we have that *f_ι_f_s_* is increasing initially around *h* ≈ 0 where *f_ι_* is flat around zero, and declining as *h → ∞* where *f_s_* is flat around zero. The home range will thus evolve to an intermediate size where foraging is optimal and the joint regulation by self-inhibition and interference is minimal, i.e., the value of *f_ι_f_s_* is at a maximum.

This equilibrium home range is defined by the optimum

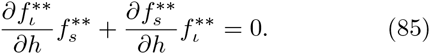

To solve, apply

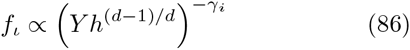

with *Y* = *nv*, *γ_i_* = *γ_l_μ*, and

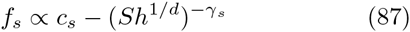

with *S* = *β/v*, as local approximations. Then,

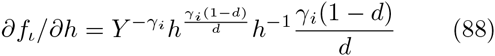

and

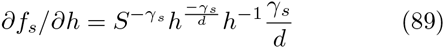

Hence, from eqn 85

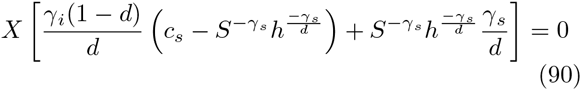

with

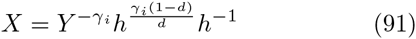

Then, divide eqn 90 with *X* on both sides, rearrange, and substitute *γ_i_* = *γ_l_μ* and *S* = *β/v* to find the optimal home range

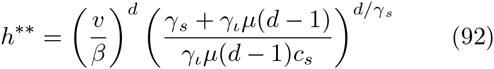

For the four traits (*v*, *β*, *n*, *h*) in the density regulation functions of interference and local resource exploitation, we find the home range (*h*) of the density regulation optimum to be density (*n*) independent, being dependent only on foraging speed (*v*) and mass specific metabolism (*β*).

If we inset eqn 92 into eqn 81 we find that the local resource exploitation at the foraging optimum 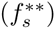 is independent of *β*, *h* and *v*, being dependent only on the other parameters of eqn 92. These are not part of the phenotype and are therefore not modified by natural selection (at least not directly), and self-inhibition at the foraging optimum is therefore expected to be invariant of the life history. The invariant local resource exploitation of optimal regulation will have an invariant derivative 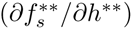, implying that eqn 85 reduces to

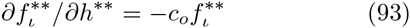

where 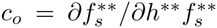 is invariant. Hence, for a given functional relation *f_l_*(*l*), we have invariant interference, and with interference in at least high-energy organisms being likely to reflect the exploitation level of the resource (*ρ_e_*), we may expect exploitation *f_e_*(*nwβ*) and density regulation as a whole *f* = *f_e_f_ι_f_s_* to be body mass invariant; generating natural selection for a trait covariance

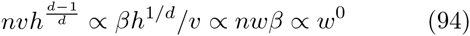

that will leave the regulation optimum invariant of the life history.

### C Replication invariance

One way of illustrating the evolutionary importance of the mass-rescaling allometries, it to focus on the average replication in a population as a function of the average mass [*pR^*^* = *f*(*w*)] for a range of potential mass-rescaling options (variation in 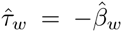). Of these it is only those with an equilibrium per generation replication rate of one (*pR^*^* = 1) that may actually be selected by natural selection.

To construct this function let 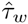 be the mass-rescaling exponent of the allometric solution (Section 3.1), let 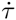 be the chosen rescaling exponent, and let the mass dependence of the juvenile period in biotic time be 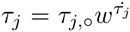, noting that this dependence in positive (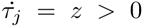, eqn 34) in the absence of mass-rescaling 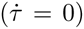, and invariant 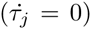 at the allometric solution (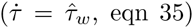, eqn 35). Hence, assuming local linearity, we find the exponent of the juvenile period 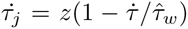 as a function of the chosen mass-rescaling exponent 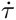. And with net energy 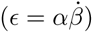 being a function of resource handling (*α*) and metabolic pace 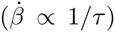, the resource handling exponent (eqn 29) is also a function of the chosen mass-rescaling with 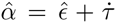. Hence, given physiological invariant survival (eqn 36), the per generation replication rate

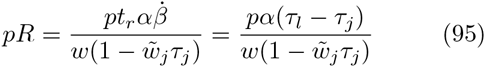

is a function of *w* given 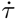 and *z*, with

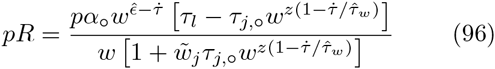

This rate is either declining or increasing with the average mass for all possible mass-rescaling options 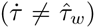 except for the allometric solution, where the per generation replication is invariant of mass (Fig. 2).

## References

Abrams P. A. Matsuda H. (1994). The evolution of traits that determine ability in competitive contests. Evol. Ecol. 8:667–686.

Anderson K. J. Jetz W. (2005). The broad-scale ecology of energy expenditure of endotherms. Ecol. Lett. 8:310–318.

Anderson W. W. (1971). Genetic equilibrium and population growth under density-regulated selection. Am. Nat. 105:489–498.

Artacho P. Nespolo R. F. (2009). Natural selection reduces energy metabolism in the garden snail, *helix aspersa* (*cornu aspersum*). Evolution 63:1044–1050.

Baltscheffsky, H., ed (1996). Origin and Evolution of Biological Energy Conversion. VCH Publishers, New York.

Banavar J. R., Damuth J., Maritan A., Rinaldo A. (2002a). Modelling universality and scaling. Nature 420:626.

Banavar J. R., Damuth J., Maritan A., Rinaldo A. (2002b). Supply-demand balance and metabolic scaling. Proc. Nat. Acad. Sci. USA 99:10506–10509.

Banavar J. R., Maritan A., Rinaldo A. (1999). Size and form in efficient transportation networks. Nature 399:130–132.

Barve A., Hosseini S. R., Martin O. C., Wagner A. (2014). Historical contingency and the gradual evolution of metabolic properties in central carbon and genomescale metabolisms. BMC Syst. Biol. 8:48.

Blum J. J. (1977). On the geometry of four-dimensions and the relationship between metabolism and body mass. J. theor. Biol. 64:599–601.

Bonner J. T. (1965). Size and cycle. Princeton University Press, Princeton.

Boratynski Z., Koskela E., Mappes T., Oksanen T. A. (2010). Sex-specific selection on energy metabolism selection coefficients for winter survival. J. Evol. Biol. 23:1969–1978.

Bozinovic F., Muñoz J. L. P., Croz-Neto A. P. (2007). In-traspecfic variability in the basal metabolic rate: testing the food habits hypothesis. Physiolo Biochem Zool 80:452–460.

Bozinovic F., Rojas J. M., Broitman B. R., Vasquez R. A. (2009). Basal metabolism is correlated with habitat productivity among populations of degus (*octodon degus*). Comp. Bioch. Physiol. A. 152:560–564.

Brody S. (1945). Bioenergetics and growth. Hafner, New York.

Brown J. H., Gillooly A. P., Allen V. M., Savage G. B. (2004). Towards a metabolic theory of ecology. Ecology 85:1771–1789.

Brown J. H. Maurer B. A. (1986). Body size, ecological dominance and cope’s rule. Nature 324:248–250.

Bulmer M. (1994). Theoretical evolutionary ecology. Sinauer Associates Publishers, Massachusetts.

Calder W. A. I. (1984). Size, function, and life history. Harvard University Press, Cambridge.

Capellini I., Venditti C., Barton R. A. (2010). Phylogeny and metabolic scaling in mammals. Ecology 91:2783–2793.

Careau V., Morand-Ferron J., Thomas D. (2007). Basal metabolic rate of canidae from hot deserts to cold arctic climates. J. Mamm. 88:394–400.

Charlesworth B. (1971). Selection in density-regulated populations. Ecology 52:469–474.

Charlesworth B. (1980). Evolution in age-structured populations. Cambridge University Press, Cambridge.

Charlesworth B. (1994). Evolution in age-structured populations. 2nd edn. Cambridge University Press, Cambridge.

Chela-Flores, J., Chadha, M., Negrón-Mendoza, A., & Oshima, T., eds (1995). Chemical Evolution: Self-Organization of the Macromolecules of Life. A. Deepak Publishing, Hampton.

Clarke B. (1972). Density-dependent selection. Am. Nat. 106:1–13.

Cunchilos C. Lecointre G. (2003). Evolution of amino acid metabolism inferred through cladistic analysis. J. Biol. Chemi. 278:47960–47970.

Currie D. J. Fritz J. T. (1993). Global patterns of animal abundance and species energy use. Oikos 67:56–68.

Damuth J. (1981). Population density and body size in mammals. Nature 290:699–700.

Damuth J. (1987). Interspecific allometry of population density in mammals and other animals: the independence of body mass and population energy-use. Biol. J. Linnean Society. 31:193–246.

Darveau C.-A., Suarez K. S., Andrews R. D., Hochachka P. W. (2002). A general model for ontogenetic growth. Nature 417:166–170.

Davison J. (1955). Body weight, cell surface and metabolic rate in anuran Amphibia. Biol. Bull. 109:407–419.

deMagalhães J. P., Costa J., Church G. M. (2007). An analysis of the relationship between metabolism, developmental schedules, and longevity using phylogenetic independent contrasts. J. Geron. Biol. Sci. 62A:149–160.

Deeds E. J. (2011). Curvature in metabolic scaling: A reply to mackay. J. theor. Biol. 280:197–198.

DeLong J. P., Okie J. G., Moses M. E., Sibly R. M., Brown J. H. (2010). Shifts in metabolic scaling, production, and efficiency across major evolutionary transitions of life. Proc. Nat. Acad. Sci. 107:12941–12945.

Demetrius L. (2003). Quantum statistics and allometric scaling of organisms. Physica 322:477–490.

Demetrius L. (2006). The origin of allometric scaling laws in biology. J. theor. Biol. 243:455–467.

Dercole F., Ferriére R., Rinaldi S. (2002). Ecological bistability and evolutionary reversals under asymmetrical competition. Evolution 56:1081–1090.

Dieckmann U. Metz J. A. J. (2006). Surprising evolutionary predictions from enhanced ecological realism. Theor. Pop. Biol. 69:263–281.

Dodds P. S., Rothman D. H., Weitz J. S. (2001). Reexamination of the “3/4-law” of metabolism. J. theor. Biol. 209:9–27.

Dreyer O. Puzio R. (2001). Allometric scaling in animals and plants. J. Math. Biol. 43:144–156.

Duncan R. P., Forsythe D. M., Hone J. (2007). testing the metabolic theory of ecology: allometric scaling exponents in mammals. Ecology 88:324–333.

Ehnes R. B., Rall B., Brose U. (2011). Phylogenetic grouping, curvature and metabolic scaling in terrestrial invertebrates. Ecol. Lett. 14:993–1000.

Fenchel T. (1974). Intrinsic rate of natural increase: The relationship with body size. Oecologia 14:317–326.

Fernando C. Rowe J. (2007). Natural selection in chemical evolution. J. theor. Biol. 247:152–167.

Fernando C. Rowe J. (2008). The origin of autonomous agents by natural selection. Bio Systems 2:355–373.

Ferry J. G. House C. H. (2006). The stepwise evolution of early life driven by energy conservation. Mol. Biol. Evol. 23:1286–1292.

Fisher R. A. (1930). The genetical theory of natural selection. Clarendon, Oxford.

Fry I. (2011). The role of natural selection in the origin of life. Origins Life Evol. Bios. 41:3–16.

Fujiwara N. (2003). Origin of the scaling rule for fundamental living organisms based on thermodynamics. BioSys. 70:1–7.

Garland T. (1983). Scaling the ecological cost of transport to body mass in terrestrial mammals. Am. Nat. 121:571–587.

Genoud M. (2002). Comparative studies of basal rate of metabolism in primates. Evol. Anthro. (Suppl) 1:108–111.

Gillooly J. F., Brown J. H., West G. B., Savage V. M., Charnov E. L. (2001). Effects of size and temperature on metabolic rate. Science 293:2248–2251.

Ginzburg L. Damuth J. (2008). The space-lifetime hypothesis: viewing organisms in four dimensions, literally. Am. Nat. 171:125–131.

Glazier D. S. (2005). Beyond the ‘3/4-power law’: variation in the intra- and interspecific scaling of metabolic rate in animals. Biol. Rev. 80:611–662.

Glazier D. S. (2008). Effects of metabolic level on the body size scaling of metabolic rate in birds and mammals. Proc. R. Soc. B. 22:821–828.

Glazier D. S. (2009). Metabolic level and size scaling of rates of respiration and growth in unicellular organisms. Funct. Ecol. 23:963–968.

Glazier D. S. (2010). A unifying explanation for diverse metabolic scaling in animals and plants. Biol. Rev. 85:111–138.

Glazier D. S. (2015). Is metabolic rate a universal ‘pacemaker’ for biological processes? Biol. Rev. 90:377–407.

Gyllenberg M. Parvinen K. (2001). Necessary and sufficient conditions for evolutionary suicide. Bull. Math. Biol. 63:981–993.

Haldane J. B. S. (1954). The origins of life. New Biol. 16:12–27.

Heino M., Metz J. A. J., Kaitala V. (1998). The enigma of frequency-dependent selection. Trends Ecol. Evol. 13:367–370.

Hill A. V. (1950). the dimensions of animals and their muscular dynamics. Science of Computer Programming 38:209–230.

Horowitz N. H. (1945). On the evolution of biochemical syntheses. Proc. Nat. Acad. Sci. 31:153–157.

Humphries M. M. McCann K. S. (2014). Metabolic constraints and currencies in animal ecology. metabolic ecology. J. Anim. Ecol. 83:7–19.

Isaac N. J. B. Carbone C. (2010). Why are metabolic scaling exponents so controversial? quantifying variance and testing hypotheses. Ecol. Lett. 13:728–735.

Jetz W., Freckleton R. P., McKechnie A. E. (2007). Environment, migratory tendency, phylogeny and basal metabolic rate in birds. PLOS One 3:e3261.

Kabat A. P., Blackburn T. M., McKechnie A. E., Butler P. J. (2008). Phylogenetic analysis of the allometric scaling of therapeutic regimes for birds. J. Zool. (London) 275:359–367.

Kiørboe T. Hirst A. G. (2014). Shifts in mass scaling of respiration, feeding, and growth rates across life-form transitions in marine pelagic organisms. Am. Nat. 183:E118–E130.

Kleiber M. (1932). Body and size and metabolism. Hilgardia 6:315–353.

Kolokotrones T., Savage V., Deeds E. J., Fontana W. (2010). Curvature in metabolic scaling. Nature 464:753–756.

Kooijman S. A. L. M. (2000). Dynamic energy and mass budgets in biological systems. Cambridge University Press, Cambridge.

Kozlowski J., Konarzewski M., Gawelczyk A. T. (2003a). Cell size as a link between noncoding DNA and metabolic rate scaling. Proc. Nat. Acad. Sci. USA 100:14080–14085.

Kozlowski J., Konarzewski M., Gawelczyk A. T. (2003b). Intraspecific body size optimization produce interspecific allometries. In: Blackburn T. M. Gaston K. J. (eds). Macroecology: Concepts and consequences: Blackwell, Malden, Massachusetts, pp 299-320

Kozlowski J. Weiner J. (1997). Interspecific allometries are by-products of body size optimization. Am. Nat. 149:352–380.

Lotka A. J. (1922). Contribution to the energetics of evolution. Proc. Nat. Acad. Sci. 8:147–151.

Lovegrove B. G. (2000). The zoogeography of mammalian basal metabolic rate. Am. Nat. 156:201–219.

Lovegrove B. G. (2003). The influence of climate on the basal metabolic rate of small mammals: a slow-fast metabolic continuum. J. Comp. Physiol. B. 173:87–112.

MacArthur R. H. (1962). Some generalized theorems of natural selection. Proc. Nat. Acad. Sci. USA 46:1893–1897.

MacKay N. J. (2011). Mass scale and curvature in metabolic scaling. J. theor. Biol. 280:194–196.

Makarieva A., Gorshkov V. G., Li B.-L. (2003). A note on metabolic rate dependence on body size in plants and animals. J. theor. Biol. 221:301–307.

Makarieva A. M., Gorshkov V. G., Bai-Lian L. (2005). Energetics of the smallest: do bacteria breathe at the same rate as whales. Proc. R. Soc. B. 272:2219–2224.

Makarieva A. M., Gorshkov V. G., Li B., Chown S. L., Reich P. B., Gavrilov V. M. (2008). Mean mass-specific metabolic rates are strikingly similar across life’s major domains: Evidence for life’s metabolic optimum. Proc. Nat. Acad. Sci. 105:16994–16999.

Marakushev S. A. Belonogova O. V. (2013). The origin of ancestral bacterial metabolism. Paleo. J. 47:1001–1010.

Maynard Smith J. Szathmáry E. (1995). The major transitions in evolution. W.H. Freeman Spektrum, Oxford.

McMahon T. A. (1973). Size and shape in biology. Science 179:1201–1204.

McNab B. K. (1988). Complications unherent in scaling the basal rate of metabolism in mammals. Quart. Rev. Biol. 63:25–54.

McNab B. K. (2003). Standard energetics of phyllostomid bats: the inadequacies of phylogenetic-contrast analyses. Comp. Bioch. Physiol. A. 135:357–368.

McNab B. K. (2008). An analysis of the factors that influence the level and scaling of mammalian bmr. Comp. Bioch. Physiol. A 151:5–28.

McNab B. K. Morrison P. (1963). Body temperature and metabolism in subspecies of *peromyscus* from arid and mesic environments. Ecol. Monogr. 33:63–82.

Melendez-Hevia E., Montero-Gomez N., Montero F. (2008). From prebiotic chemistry to cellular metabolism the chemical evolution of metabolism before darwinian natural selection. J. theor. Biol. 252:505–519.

Metz J. A. J., Mylius S. D., Diekmann O. (1996). When does evolution optimize? On the relation between types of density dependence and evolutionary stable life history parameters. IIASA WP:96–04.

Miller S. L. (1953). A production of amino acids under possible primitive Earth conditions. Science 117:528–529.

Mueller P. Diamond J. (2001). Metabolic rate and environmental productivity: well-provisioned animals evolved to run and idle fast. Proc. Nat. Acad. Sci. 98:12551–12554.

Muñoz-Garcia A. Williams J. B. (2005). Basal metabolic rate in carnivores is associated with diet after controlling for phylogeny. Physiolo Biochem Zool 78:1039–1056.

Mylius S. D. Diekmann O. (1995). On evolutionarily stable life histories, optimization and the need to be specific about density dependence. Oikos 74:218–224.

Nagy K. A. (2005). Field metabolic rate and body zise. J. Exp. Biol. 208:1621–1625.

Nee S., Read A. F., Greenwood J. J. D., Harvey P. H. (1991). The relationship between abundance and body size in british birds. Nature 351:312–313.

Niven J. E. Scharlemann J. P. W. (2005). Do insect metabolic rates at rest and during flight scale with body mass? Biol. Lett. 1:346–249.

Nowak R. M. (1991). Walker’s mammals of the world volume I–II. 5th edn. The Johns Hopkins University Press, Baltimore.

Oparin, A. I. & Clark, F., eds (1959). The origin of life on Earth. Pergamon Press, New York.

Padfield D., Yvon-Durocher G., Buckling A., Jennings S., Yvon-Durocher G. (2016). Rapid evolution of metabolic traits explains thermal adaptation in phytoplankton. Ecol. Lett. 19:133–142.

Parry G. D. (1981). The meanings of r- and k-selection. Oecologia 48:260–264.

Patterson M. R. (1992). A mass transfer explanation of metabolic scaling relations in some aquatic invertebrates and algae. Science 255:1421–1423.

Pawar S., Dell A. I., Savage V. M. (2012). Dimensionality of consumer search space drives trophic interaction strengths. Nature 486:485–489.

Pearl R. (1928). The rate of living. Alfred A. Knopf, New York.

Peters R. H. (1983). The ecological implication of body size. Cambridge University Press, Cambridge.

Pianka E. R. (1970). On r- and k-selection. Am. Nat. 104:592–596.

Ponnamperuma, C. & Chela-Flores, J., eds (1993). Chemical Evolution: Origin of Life. A. Deepak Publishing, Hampton.

Rau A. R. P. (2002). Biological scaling and physics. J. Biosci. 27:475–478.

Robertson A. (1968). The spectrum of genetic variation. In: Lewontin R. C. (ed). Population Biology and Evolution: Syracuse University Press, New York, pp 5–16.

Roff D. A. (1992). The evolution of life histories. Theory and analysis. University of Chicago Press, New York.

Roff D. A. (2002). Life history evolution. Sinauer Associates, Inc., Massachusetts.

Roughgarden J. (1971). Density-dependent natural selection. Ecology 5:453–468.

Santillán M. (2003). Allometric scaling law in a simple oxygen exchanging network: possible implications on the biological allometric scaling laws. J. theor. Biol. 223:249–257.

Savage V. M., Gillooly J. F., Wooduff W. H., West G. B., Allen A. P., Enquist B. J., Brown J. H. (2004). The predominance of quarter-power scaling in biology. Funct. Ecol. 18:257–282.

Schoener T. W. (1968). Sizes of feeding territories among birds. Ecology 49:123–131.

Sibly R. M., Brown J. I., Kodric-Brown A. (2012). Metabolic ecology: A scaling approach. John Wiley & Sons, Chichester.

Sibly R. M. Calow P. (1986). Physiological ecology of animals. Blackwell, Oxford.

Sieg A. E., O’Connor M. P., McNair J. N., Grant B. W., Agosta S. J., Dunham A. E. (2009). Mammalian metabolic allometry: do intraspecific variation, phylogeny, and regression models matter? Am. Nat. 175:720–733.

Smith C. C. Fretwell S. D. (1974). The optimal balance between size and number of offspring. Am. Nat. 108:499–506.

Stahl W. R. (1962). Similarity and dimensional methods in biology. Science 137:205–212.

Stearns S. C. (197). The evolution of life-history traits: A critique of the theory and a review of the data. Ann. Rev. Ecol. Syst. 8:145–171.

Stearns S. C. (1976). Life-history tactics: A review of the ideas. Quart. Rev. Biol. 51:3–47.

Stearns S. C. (1992). The evolution of life histories. Oxford University Press, Oxford.

Stearns S. C. Hoekstra R. F. (2000). Evolution: an introduction. Oxford University Press, Oxford.

Taylor P. D. (1996). The selection differential in quantitative genetics and ess models. Evolution 50:2106–2110.

Turner F. B., Jennrich R. I., Weintraub J. D. (1969). Home range and body size of lizards. Ecology 50:1076–1081.

Versteegh M. A., Schwabl I., Jaquier S., Tieleman B. I. (2012). Do immunological, endocrine and metabolic traits fall on a single pace-of-life axis? covariation and constraints among physiological systems. J. Evol. Biol. 25:1864–1876.

Weibel E. R., Bacigalupe L. D., Schmitt B., Hoppeler H. (2004). Allometric scaling of maximal metabolic rate in mammals: muscle aerobic capacity as determinant factor. Resp. Physiol. Neurobiol. 140:115–132.

West G. B., Brown J. H., Enquist B. J. (1997). A general model for the origin of allometric scaling laws in biology. Science 276:122–126.

West G. B., Brown J. H., Enquist B. J. (1999a). A general model for the structure and allometry of plant vascular systems. Nature 400:664–667.

West G. B., Brown J. H., Enquist B. J. (1999b). The fourth dimension of life: Fractal geometry and allometric scaling of organisms. Science 284:1677–1679.

White C. R., Blackburn T. M., Martin G. R., Butler P. J. (2007). Basal metabolic rate of birds is associated with habitat temperature and precipitation, not primary productivity. Proc. R. Soc. B. 274:287–293.

White C. R., Blackburn T. M., Seymour R. S. (2009). Phylogenetically informed analysis of the allometry of mammalian basal metabolic rate supports neither geometric nor quarter-power scaling. Evolution 63:2658–2667.

White C. R., Cassey P., Blackburn T. M. (2007). Allometric exponents do not support a universal metabolic allometry. Ecology 88:315–323.

White C. R. Kearney M. R. (2013). Determinants of interspecific variation in basal metabolic rate. J. Comp. Physiol. B. 183:1–26.

Wikelski M., Spinnery L., Schelsky W., Scheuerlein A., Gwinner E. (2003). Slow pace of life in tropical sedentary birds: a common-garden experiment on four stonechat populations from different latitudes. Proc. R. Soc. B. 270:2383–2388.

Withers P. C., Cooper C. E., Larcombe A. N. (2006). Enviromental correlates of physiological variables in marsupials. Physiolo Biochem Zool 79:437–453.

Witting L. (1995). The body mass allometries as evolutionarily determined by the foraging of mobile organisms. J. theor. Biol. 177:129–137.

Witting L. (1997). A general theory of evolution. By means of selection by density dependent competitive interactions. Peregrine Publisher, ?Arhus, 330 pp, URL http://mrLife.org.

Witting L. (1998). Body mass allometries caused by physiological or ecological constraints? Trends Ecol. Evol. 13:25.

Witting L. (2000). Interference competition set limits to the fundamental theorem of natural selection. Acta Biotheor. 48:107–120.

Witting L. (2002). Two contrasting interpretations of Fisher’s fundamental theorem of natural selection. Comm. Theor. Biol. 7:1–10.

Witting L. (2003). Major life-history transitions by deterministic directional natural selection. J. theor. Biol. 225:389–406.

Witting L. (2008). Inevitable evolution: back to *The Origin* and beyond the 20th Century paradigm of contingent evolution by historical natural selection. Biol. Rev. 83:259–294.

Witting L. (2016a). The natural selection of metabolism and mass selects lifeforms from viruses to multicellular animals. bioRxiv http://dx.doi.org/10.1101/087650.

Witting L. (2016b). The natural selection of metabolism bends body mass evolution in time. bioRxiv http://dx.doi.org/10.1101/088997.

Witting L. (2016c). The natural selection of metabolism explains curvature in allometric scaling. bioRxiv http://dx.doi.org/10.1101/090191.

